# New insight in cyclic monoterpenoids mechanism of action: an in silico approach

**DOI:** 10.1101/2024.06.05.597591

**Authors:** Silvia Pezzola, Federica Sabuzi, Pierluca Galloni, Valeria Conte, Mariano Venanzi, Gianfranco Bocchinfuso

## Abstract

Clarifying the mechanism of action of natural terpenoids is challenging. Further, their efficacy is inspiring in developing new antimycotic agents. Among all, thymol, carvacrol and thymyl acetate are largely scrutinized, while the new brominate thymol, namely bromothymol (4-bromo-2-isopropyl-5-methylphenol), needs deeper investigation. Here its antimycotic efficacy was evaluated and, in parallel, a careful in silico investigation of the mechanism of action was proposed. In vivo experiments, on species of acclaimed resistance, demonstrated that bromothymol had a Minimum Inhibitory Concentration (MIC) equal ∼40 μg/ml, 6 times more active than thymol. Partition coefficient (LogP) in heptane, determined through density functional theory (DFT), and Molecular Dynamics (MD) simulations, based on a Minimum Bias Approach, in the presence of neutral bilayers, indicated that bromothymol inserts into cellular membrane, such as thymol, carvacrol, and Thymyl acetate. Monoterpenoids bearing the hydroxyl group induces a shrinkage of the membrane thickness, while only thymol affected membrane density of the leaflets in which it inserted. Thymol, carvacrol, and bromothymol interacted with the polar head of the lipids causing an electrostatic imbalance into the membrane, justifying their biological activity. For the first time a detailed in silico characterization on the mechanism of these compounds is afforded, returning a coherent and clear picture of their mechanism of action.

## 1. Introduction

The spread of new moulds in food and crops, often resistant to ordinary treatments, has evidenced the need to develop innovative phyto-molecules [1-4]. In this context, natural monoterpenoids, and their tailored substituents, have been recently re-discovered, being inspired by traditional medicine and plants defence mechanism [5-8]. Among the essential oils of the officinalis plants [5], thymol (2-isopropyl-5-metilphenol), carvacrol (5-isopropyl-2-metilphenol), geraniol (3,7-dimetil-2,6-ottadien-1-ol), and eugenol (a-allyl-2-metoxyphenol), are probably the best characterized from the chemical and physicochemical point of views [9]. New insight reveals that thymol and carvacrol, acting on planktonic cells, prevented and eradicated biofilm formation, disrupting pathogens cells, [10-11], strengthening the interest in understanding their pathway of action. A large body of literature assesses that they penetrate the cell membrane, inducing its morphological collapse and, in the end, causing its disruption [12]. Properly, thymol increased the curvature of the membrane surface, inducing massive changes in its properties, such as a decrease in elasticity and an increase in fluidity [10, 13]. Furthermore, epi-fluorescent and NMR studies, performed on a Langmuir film as a model membrane, suggested that thymol, propofol (2,6-diisopropylhenols), carvacrol, and eugenol interact with the apical region of membranes. Experimentally, it was found that they located in the choline region of the lipids, diminished the repulsion among phospholipids head-groups, and formed hydrogen bonds with the glycerol and/or phosphate moiety [14-15]. Other investigations hypothesized a similar pattern also behaviour on bacteria membrane, without leading to any unquestionable mechanism [9, 13]. Elsewhere, the exploitation of Langmuir monolayers confirmed that thymol altered the smoothness and the density of the system, causing a significant expansion [16]. Thus, the umbrella sampling on dipalmitoyl-phosphatidylcholine monolayer was performed in predicting the pose of thymol, concluding that it is placed in the hydrophobic region of the membrane [16]. However, authors did not detail a mechanism of action. In vivo investigation, on Botrytis cinerea, a fungus causing cereal stem and fruit rot, demonstrated that carvacrol had less efficacy than thymol in inducing hyphae, and/or spore death through their shrinkage [17]. Therefore, the authors claimed that both monoterpenoids prompted a sudden reduction of the media pH, a release of the K+ from cellular environment, an increase of extracellular conductivity, and a change in membrane permeability [17]. Wanting to explain the lower efficacy of carvacrol, umbrella sampling was exploited, but no relevant result was highlighted [14, 15, 18]. Overall, the data suggest that thymol, carvacrol and their tailored analogues, strictly interact with the cellular membrane without allowing the development of any resistance mechanism into host cells, and, in parallel, inhibit/prevent biofilm formation. All these features make these tempting compounds to be exploited as active principles in pharmaceutical/disinfectant fields [5,10, 13].

Computational techniques, such as QM and/or MD, are widely used to predict and characterize molecule-molecule interactions. DFT allows computing compound reactive moieties (i.e. EPM), their stability (i.e. FMO), and their hydrophobicity towards hydrophilic/hydrophobic environment (i.e. LogP) [19-22]. By restricting the field to the interaction with bilayer, between others, Two MD computational approaches can be used to predict the more stable configurations of the system: i) the umbrella sampling, accomplished by the calculation of the potential mean force (PMF), and ii) the Minimum Bias Approach (MBA) [16, 23]. In the PMF approach, compounds are pushed through a preassembled double layer membrane. They pass through several virtual poses, until reaching “putative” energetic minimum. In the MBA, instead, simulations started form a chaotic mixture of lipids, water, ions and the investigated compound. Notwithstanding the lower computational cost with respect to the US-PMF methods, MBA is able to individuate the most stable configurations of the system [23] and it is useful to determine and clarify the interaction of small peptide or organic compounds with membranes at the atomic level [24-30].

In the present work, the antifungal activity of bromothymol, namely the brominated tailored analogue of thymol in position 4, was compared to that of thymol, exploiting a selected panel of potential pathogen yeasts and moulds. Moreover, a detailed in silico analysis was performed to clarify bromothymol mechanism of action in comparison to that of thymol carvacrol and thymyl acetate. These compounds were selected as internal standard of the suggested approach, due to the large amount of information on their activity. DFT was exploited in calculating the EPM, the FMO, and the LogP of each compound. POPC was selected as model of a simplified eukaryote membrane in MBA studies. Relative placement of each compound, hydrogen bonds formation, effects on membrane density and thickness, as well as lipid modifications, were also investigated.

## 2. Results and Discussion

### 2.1 Antifungal activity

Bromothymol in vivo efficacy was investigated on potentially pathogen moulds and yeasts, selected because they affect commercial goods, as well as human and animal health [1-3]. Contrary to carvacrol, thymol, and its acetylated form, thymyl acetate [9, 16, 31, 32-33], little is known about bromothymol [34]. Bromothymol MIC was compared to the thymol one, the reference compound in commercial application. MIC was determined analysing thymol and bromothymol concentrations from 0 (control) to 900 μg/ml (Figure 1 and Table S1).

**Figure 1.**
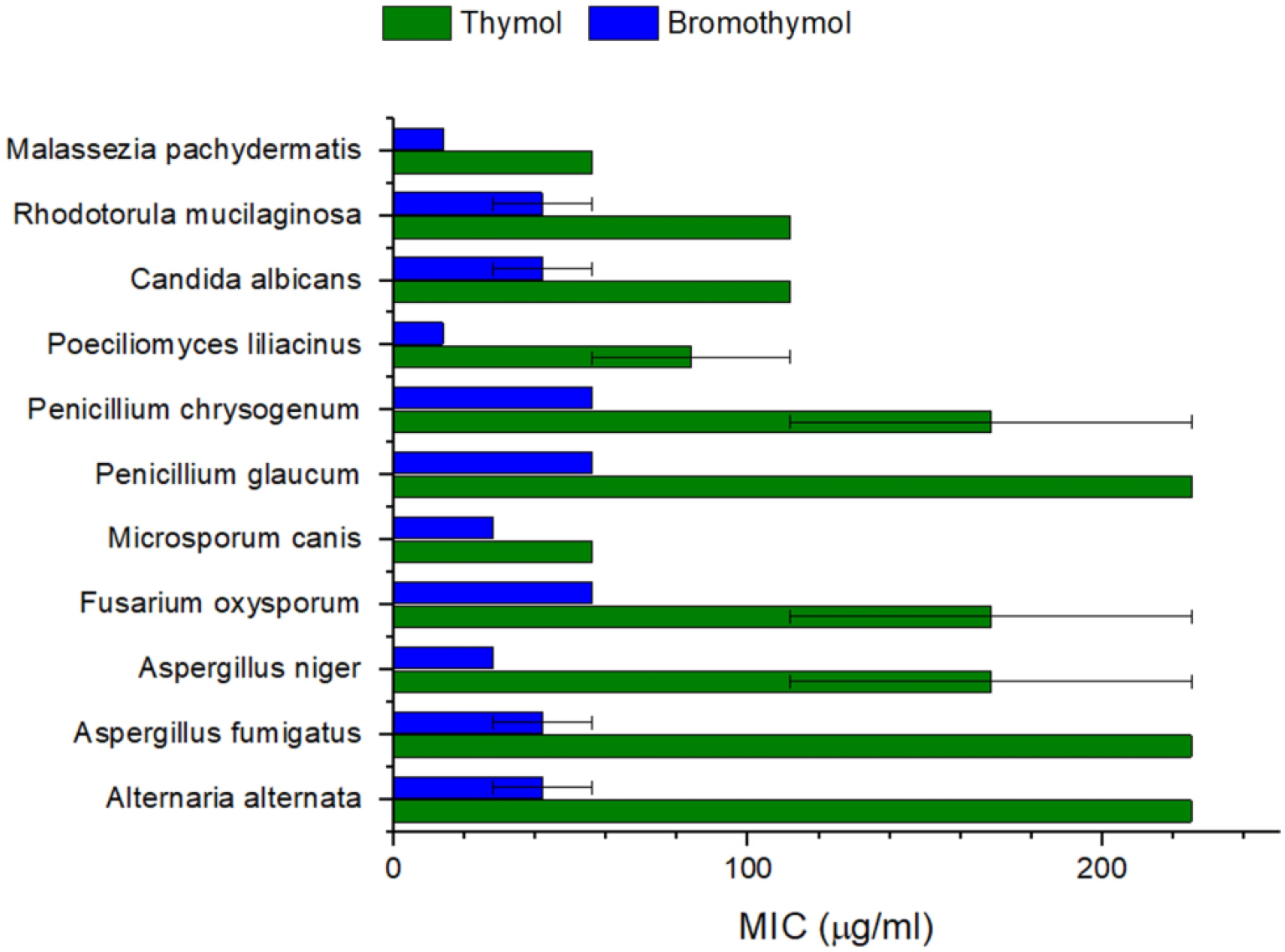
MIC determination of thymol (green) and bromothymol (blue) on a selected panel of yeast and moulds.

Noteworthy, the brominated form was almost four times more active than thymol on the considered strains. For instance, bromothymol MIC was at least two times lower than that of thymol in the difficult-to-treat M. canis (J Fungi (Basel). 2018 Sep; 4(3): 107). Remarkably, the functionalized monoterpenoid was up to 5 time more active than the natural one, on A. fumigatus, A. niger, F. oxysporum, P. glaucum, and P. chrysogenum. Namely, Figure 1 and TableS1. In Alternaria alternata, an opportunistic pathogen [1-3], bromothymol MIC was up to six time lower than the thymol one, TableS1.Eventually, bromothymol was four time more active also against M. pachydermatis and P. liliacinus, which were well-known for their resistance to most of the commercially available antimycotic compounds, TableS1.

### 2.2 The mechanism of action

A piece of evidence suggests that the thymol derivatives act by permeabilizing the pathogen membranes. This phenomenon is the combination of several effects, such as hydrophobicity, electrostatic interactions, ionic strength of the solution, dynamics of the membrane and how compound acts on it. Altogether, these features affect the final activity of each specific compound. The overall complexity can be reduced by considering two simple facts: i) the higher the tendency of the compound to bind with the membrane, the higher its activity; ii) at an equal affinity for the membrane, the higher the perturbation induced to the bilayer the higher the activity. Either of these general characteristics can be based on the different activities of distinctive molecules. Computational techniques exploitation is pivotal in clarifying the structural reason of the observed higher activity of the bromothymol with respect to the thymol. Further, it is a needed step in developing new and more powerful molecules. Thus, bromothymol, thymol, carvacrol, and thymil-acetate ability to bind membrane was investigate through quantum mechanical (QM) prediction of the LogP (paragraph 2.2.1) and the effect on the membrane dynamics was evaluated by means of MD simulations in the presence of POPC bilayers (paragraph 2.2.2). Finally, the potential reactivity of each compound was also investigated by means of DFT calculations (paragraph 2.2.3), wanting to examinate the possibility of exploiting these compounds in further applications requiring chemical stability (such as photo-induced perturbation of targeted membranes).

#### 2.2.1 The compounds show different tendencies to bind to the membrane

The tendency of a molecule to interact with a bilayer is affected by different factors, however, the logP represents a simple parameter for a first assessment of this behaviour [35]. Beyond this aspect, the logP evaluation is a pivotal characteristic in drug discovery and development because it influences the absorption, distribution, metabolism, excretion, and toxicity (ADMET) [36] LogP is defined as the log of the partition coefficient of a solute between octanol and water, at near infinite dilution, however, octanol can form hydrogen bonds with solutes and water, therefore, hydrocarbons/water partition is generally considered to be more appropriate in modelling the interaction with the hydrophobic core of a membrane bilayer [37]. Here, LogP in heptane/water was calculated in accordance with equation 1.

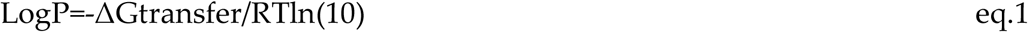

where R is the Gas constant, T is the temperature (herein 298.15K) and ΔGtransfer is the difference in Gibbs free energy (ΔG) of water and heptane, respectively [38]. Calculations were performed with B3LYP as functional, according to the Automate Topology builder (ATB) output, further exploited in MD. Solvation model base on density (SMD) was selected for its capability in well describe water and non-polar solvents, while 6-311G+(d, p) as basis set, because it accurately foresees hydrogen-bond interaction. The calculated heptane/water was compared to the available experimental ones, wanting to predict compound behaviour in lipids-like environment (Table 2).

**Table 2.** Calculation of partition coefficient in heptane/water for: thymol, carvacrol, bromothymol, and thymyl acetate. Data were obtained after geometry optimization with B3LYP/SMD/6-311G+(d,p) in water and heptane, respectively. LogPexp values were obtained from the NIH (National Istitute of Health) through PubChem [39], ND stands for Not Determined.

| Compound | LogPe<br>xp | LogPHept/<br>water |
| --- | --- | --- |
| Thymol | 3.3 | 1.68 |
| Carvacrol | 3.49 | 1.83 |
| Bromothymol | ND | 1.96 |
| Thymil-acetate | 3 | 1.52 |
| R-Square |  | 0.99 |

Computed LogP_Hept/water_ showed a similar trend with the available experimental data [39]. Bromothymol shows the highest value of computed LogP among the analysed molecules, suggesting a major affinity towards the double layer, also in agreement with its higher in vivo efficacy. Therefore, the insertion of bromothymol in the bilayer was afforded to investigate the possibility that it can produce, or not, a more dramatic perturbation of the bilayer properties with respect to the other compounds.

#### 2.2.3 How the thymol derivatives insertion perturbs the bilayer properties

To evaluate the effect on the membrane, the assembling of bilayers in the presence of one of the thymol derivatives was simulated. Selected conditions mimic low compounds concentration. Whilst the bilayer is forming, the compounds of interest can efficiently explore all the conformational space. Once formed the bilayer, the systems lost fluidity and the reached configuration is stably retained for different hundreds of nanoseconds. By replicating the protocol many times, the ensemble of the accessible configurations is sampled, and the effects on the dynamic properties of the membrane can be investigated without introducing any bias (such as coordinate reactions); for this reason, this is sometimes referred to as the Minimum Bias Approach (MBA). In this study, ten independent boxes were built for each compound, and a minimum of seven, showing a defect-free bilayer, were analysed (Sup. Info Figure S1). MBA output revealed that all the simulated compounds penetrated the bilayer, according to the predicted LogP (Table 2). To better clarify molecules interaction with the bilayer, the deepness of different functional groups was thus inspected. Especially, the alcoholic or acetylic group, potentially involved in hydrogen bonds, the isopropyl moiety, and the aromatic group involved in weak non-polar interaction were investigated [19, 31-33, 40-42]. To this end, Figure S2 reports the density profile along the axis orthogonal to the bilayer plane.

All the thymol derivatives are located close to the water/bilayer interface, slightly below the lipid polar heads, with the hydroxyl group more exposed to the solvent. In the bromothymol, the bromine, in para position with respect to the hydroxyl group, is the deeper inserted one.

The aromatic moiety is the common structural feature among the investigated molecules, thus the Radial distribution function (RDF) of the phosphate group (P8, Scheme S1) and of the ester oxygens (O14 and O16, Scheme S1) with respect to the benzene ring was calculated (Figure S3). Overall, these data show a quite similar behaviour of thymol, carvacrol and bromothymol, with well-defined peaks roughly centred at 4.5 and 6.0 Å for phosphate and 3.5 and 4.5 Å for the oxygens. Thymyl acetate loss this structured pattern, witnessing a completely different dynamics of the molecule in the bilayer, well evident by visual inspection of the trajectories. Furthermore, the absence of the hydroxyl group caused a lower retention of the molecule in the region close to the water phase. The ability of the hydroxyl group in thymol carvacrol and bromothymol to form H-bond was also investigated. Figure 2 reports the frequency of H-bond with water and with the etheric oxygen (O7, Scheme S1) during the last 10 ns of the dynamics.

**Figure 2.**
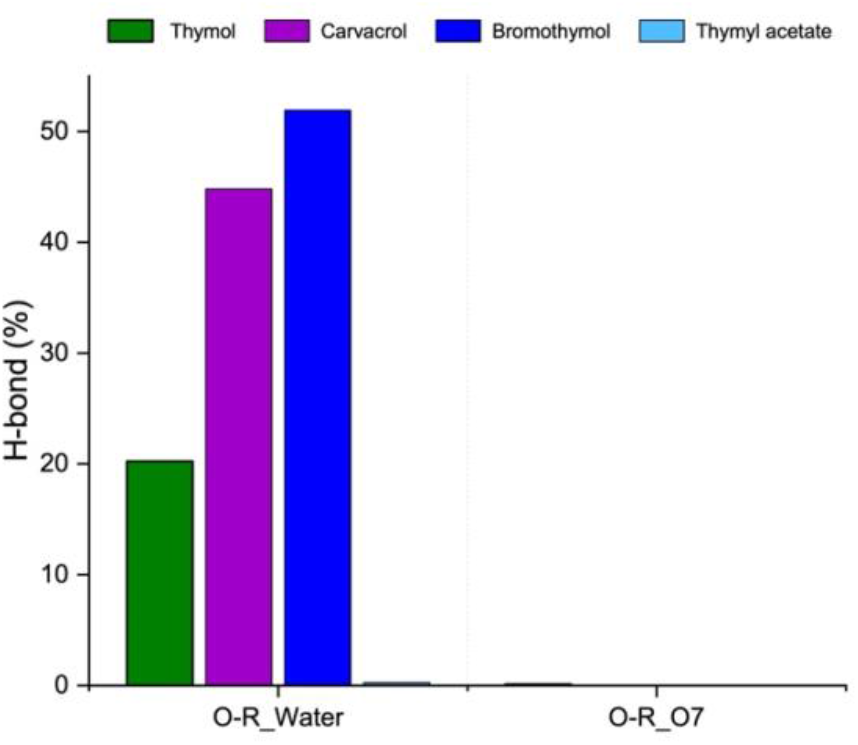
Average percentage of hydrogen bonding formation during the last 10 ns of dynamics. R: hydrogen in thymol, carvacrol and bromothymol, OR: etheric group in thymyl acetate. O7: choline oxygen of lipids

In bromothymol, hydrogen bond percentage was up to 52.0%±0.2, implying its tight interaction with the solvent and/or lipids moiety able to act as hydrogen bond donor or acceptor, in accordance with other monoterpenoids behaviour [42]. Even carvacrol, conversely from thymol, formed up to 44% of hydrogen bond with solvent. For all the compounds, weaker interactions were established with the O7, contrary to what was stated before [14-15], suggesting that the choline oxygen is not the main target of the hydrogen bond. Further, the O1 of thymol, carvacrol and bromothymol is very close to the P8, (roughly 3.4 Å), implying a more strength interaction with that group (Figure S4).

P8 is accounted as the limit between the hydrophilic-hydrophobic threshold, thus the distance O1-P8 is linked to the depth that compounds reached into the bilayer. Thymyl acetate inner depth (ID) was equal to 8.2 ±0.7 Å, meanwhile, bromothymol (ID=4.3±0.8 Å), carvacrol (ID= 4.2±0.6 Å) and thymol (ID= 4.2 ±0.5 Å) mostly remained into the hydrophilic region. This result confirms that compounds bearing a H-bond donor moiety are “trapped” into a fixed position, while Thymyl acetate, not bearing H-bond donor moiety, shows a higher mobility and a deeper penetration, according to its LogP value.

Overall, the analysed compounds are in the centred between the polar and the non-polar regions of the bilayer, potentially playing a perturbative role on the aliphatic chains and the polar heads, as well. To investigate the effect on the aliphatic chains, the order parameters in the presence or not of the thymol derivatives have been afforded (Figure 3).

**Figure 3.**
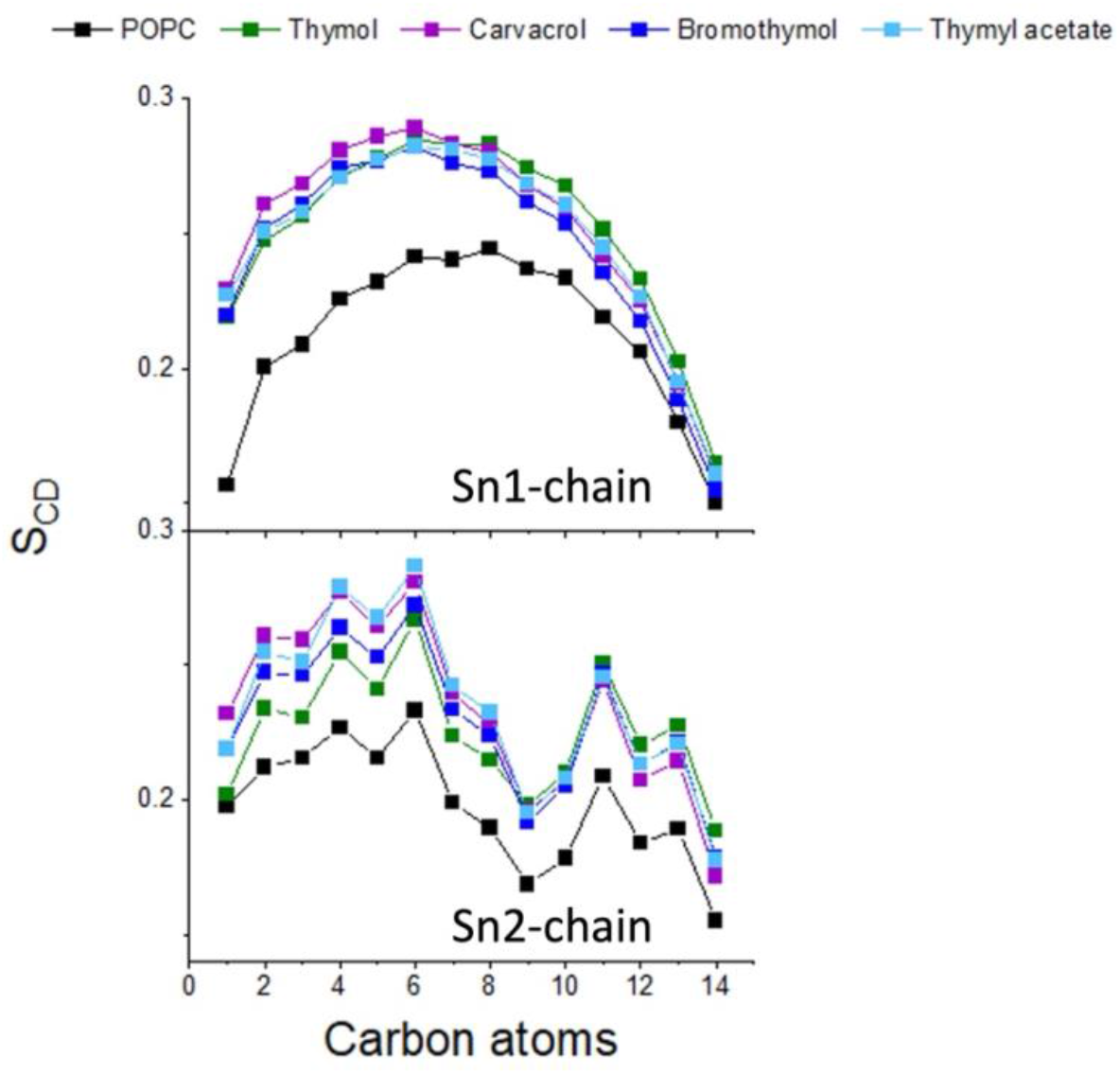
Order parameter (SCD) of lipid bilayer in the presence and in the absence of compounds. a) sn1-chain of lipid and b) sn2-chain of lipid.

Even if monoterpenoids did not induce the transition to a gel-like phase, (SCD< 0.3), [42], a huge perturbation of the palmitic and oleic chains, sn1 and sn2-chain, with respect to the POPC alone was observable. However, a peculiar behaviour of the bromothymol has not been detected; this shared behaviour did not explain differences in the biological activity of compounds.

Because it is commonly accepted that thymol plays a key role in modifying membrane morphology [10, 13, 14], the possibility in emphasizing its alteration was also afforded, Figure 4.

**Figure 4.**
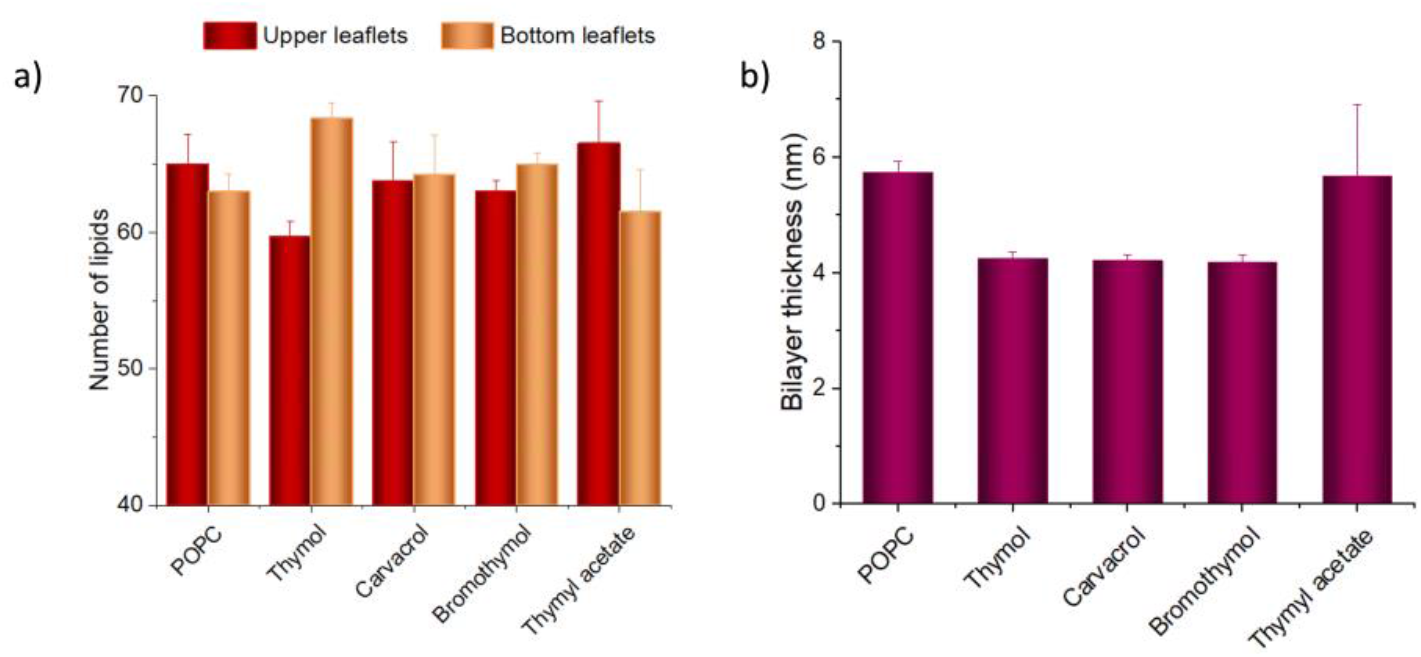
Lipids distribution between the two leaflets in the presence or not of the target compound. Panel a) Density variation, determined as the distribution of lipids between the double layer freely formed under MBA conditions., upper leaflet was chosen as the one where drugs were posed into, and further compared with the control (lipids only). Panel b): bilayer thickness variation in presence or not of different compounds.

Alteration in leaflet density was calculated as the variation in the number of lipids distribution between the leaflets in presence or not of compounds, taking advantage of the MBA performance [23]. In the control membrane, lipid equally distributed between the two leaflets (Δlipids=2± 1.2), (Figure 4, panel a). Noteworthy, only thymol induce a significant imbalance in the lipids number, (Δlipids=8±1,54), decreasing the membrane density, implying an increasing its curvature. Otherwise, all phenols derivatives bearing the -OH group induced a shrinkage in the double layer, that varied from 5 nm, in untreated condition, up to 4.2 in the presence of bromothymol, data that was only hypothesized previously [14-15, 42].

So far, MBA approach reaps a coherent picture of the mechanism of action, fulfilling the jigsaw presents by the literature. However, further analysis was compelled to clarify the structural reasons since the higher activity of the bromothymol. In the next, the effect on the dynamics features of the polar heads of the compounds bearing a hydroxyl group was investigated (i.e., thymol, carvacrol, and bromothymol). A visual inspection of the acquired trajectories suggests that two possible poses are allowed, characterized a different H bond pattern of the hydroxyl group in the molecule : i) H-bonds with the oxygens of the phosphate group (O9/O10 Scheme S1) (POSE_1); or ii) H-bonds with the oxygens belonging to the palmitoyl tail (O14/O16 Scheme S1), POSE_2, (Figure 5). These interactions are of major interest, as the activity of monoterpenoids is supposed to be mediated by these H-bonds [9, 14-15].

**Figure 5.**
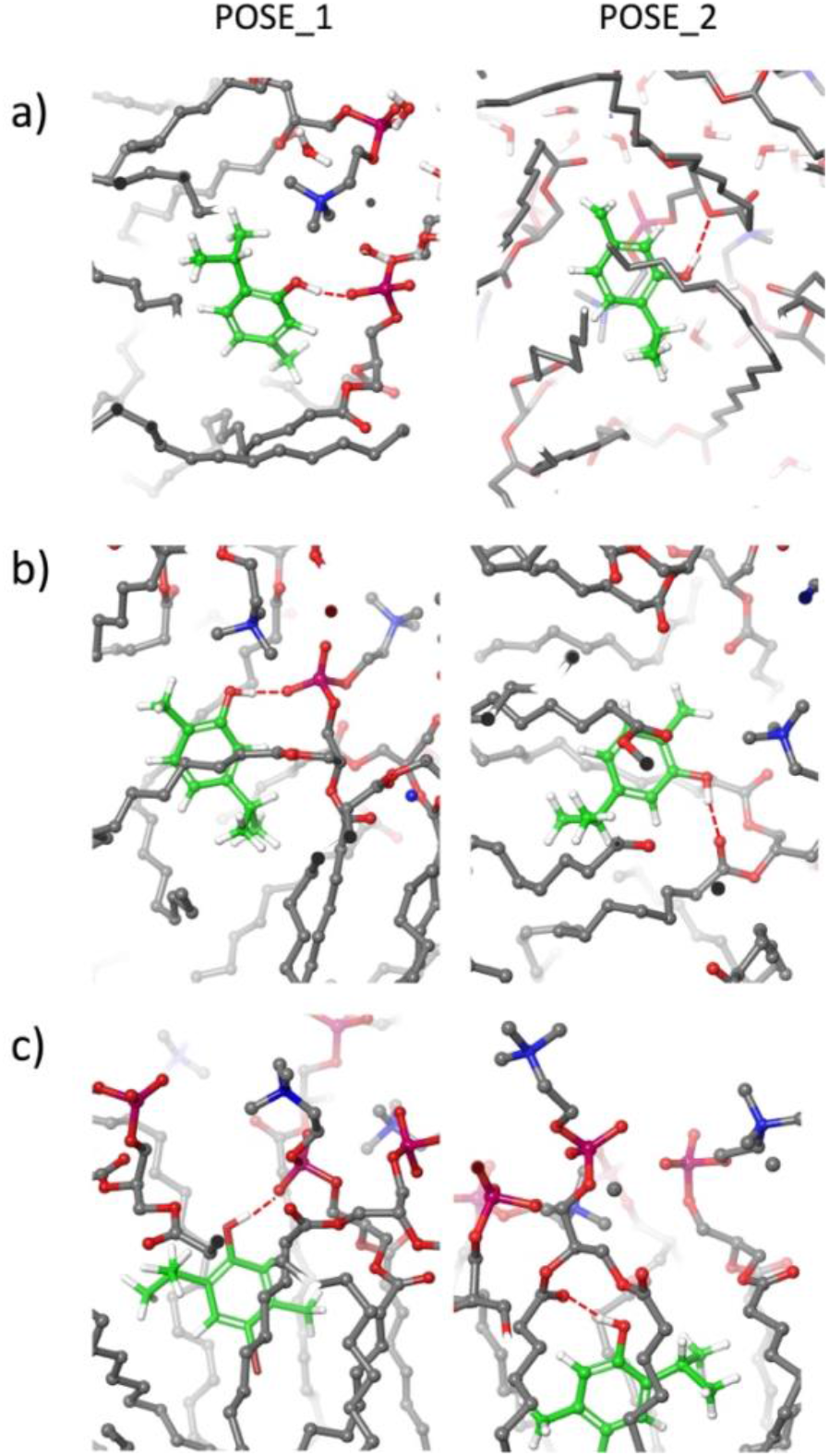
Thymol, carvacrol and bromothymol pose into the double layer. POSE_1: formation of a hydrogen bond between the oxygen linked to the phosphate and the hydrogen of the -OH group. POSE_2: formation of a hydrogen bond between the oxygen of the palmitoyl chain and the hydrogen of the -OH group. Red dotted line: hydrogen-bond. Panel: a) thymol; b) carvacrol; c) bromothymol.

When POSE_1 or POSE_2 was established, it was constant throughout the dynamics. Interestingly, the poses showed different frequencies for the three compounds (Table 3).

**Table 3.** Percentage of compounds distribution in POSE_1 and POSE_2, among the seven investigated dynamics.

| Compound | POSE_1 | POSE_2 |
| --- | --- | --- |
| Thymol | 86% | 14% |
| Carvacrol | 40% | 60% |
| Bromothymol | 60% | 40% |

Thymol assumed mainly the POSE_1 in the 86% of the dynamics, binding the O9 (see Scheme S1, for clarity), whereas, with slight differences, carvacrol and bromothymol allowed both placements. Noteworthy, these data resolve an apparent discrepancy presents in the literature, where two different poses for thymol and carvacrol were already proposed, but as alternative to each other [14, 16].

Single point energy experiment was planned to understand this frequency disparity among compound placements. Geometry optimization was performed with CAM-B3LYP, due to its capability to predict long-range interaction and hydrogen bonds, and 6-31 was selected as basis set, because of the complexity of the system. For each molecule, the difference between POSE_1 and POSE_2 (ΔE) was calculated, Table 4.

**Table 4.**
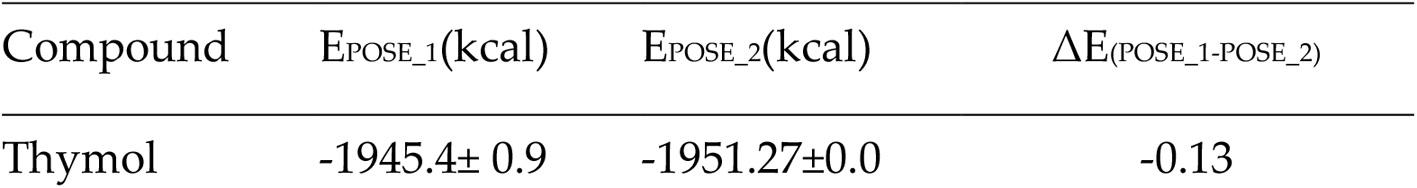
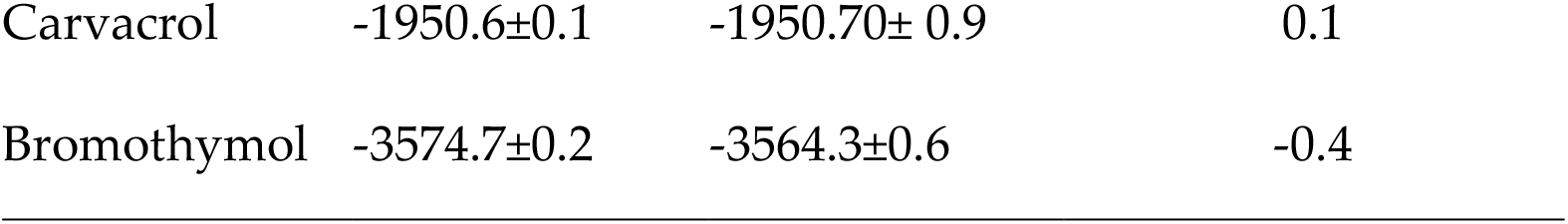
Average energy associated to POSE_1 and POSE_2 of the investigated compounds. Geometry optimization was performed with CAM-B3LYP, in vacuum, and exploiting 6-31 as basis set.

| Compound | $E_{\text{POSE}_1}(\text{kcal})$ | $E_{\text{POSE}_2}(\text{kcal})$ | $\Delta E_{(\text{POSE}_1-\text{POSE}_2)}$ |
| --- | --- | --- | --- |
| Thymol | -1945.4± 0.9 | -1951.27±0.0 | -0.13 |
| Carvacrol | -1950.6±0.1 | -1950.70± 0.9 | 0.1 |
| Bromothymol | -3574.7±0.2 | -3564.3±0.6 | -0.4 |

Data in Table 4 are in essential agreement with MD, explaining the difference in frequency output. The two different poses appear to affect the polar heads conformations. As representative of this effect, the distance between the phosphate and nitrogen atoms in the lipids was analysed (Figure 6 and table S2). 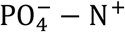 in the lipids form a dipole and this distance is pivotal for the electrostatic properties of the POPC bilayer [43].

**Figure 6.**
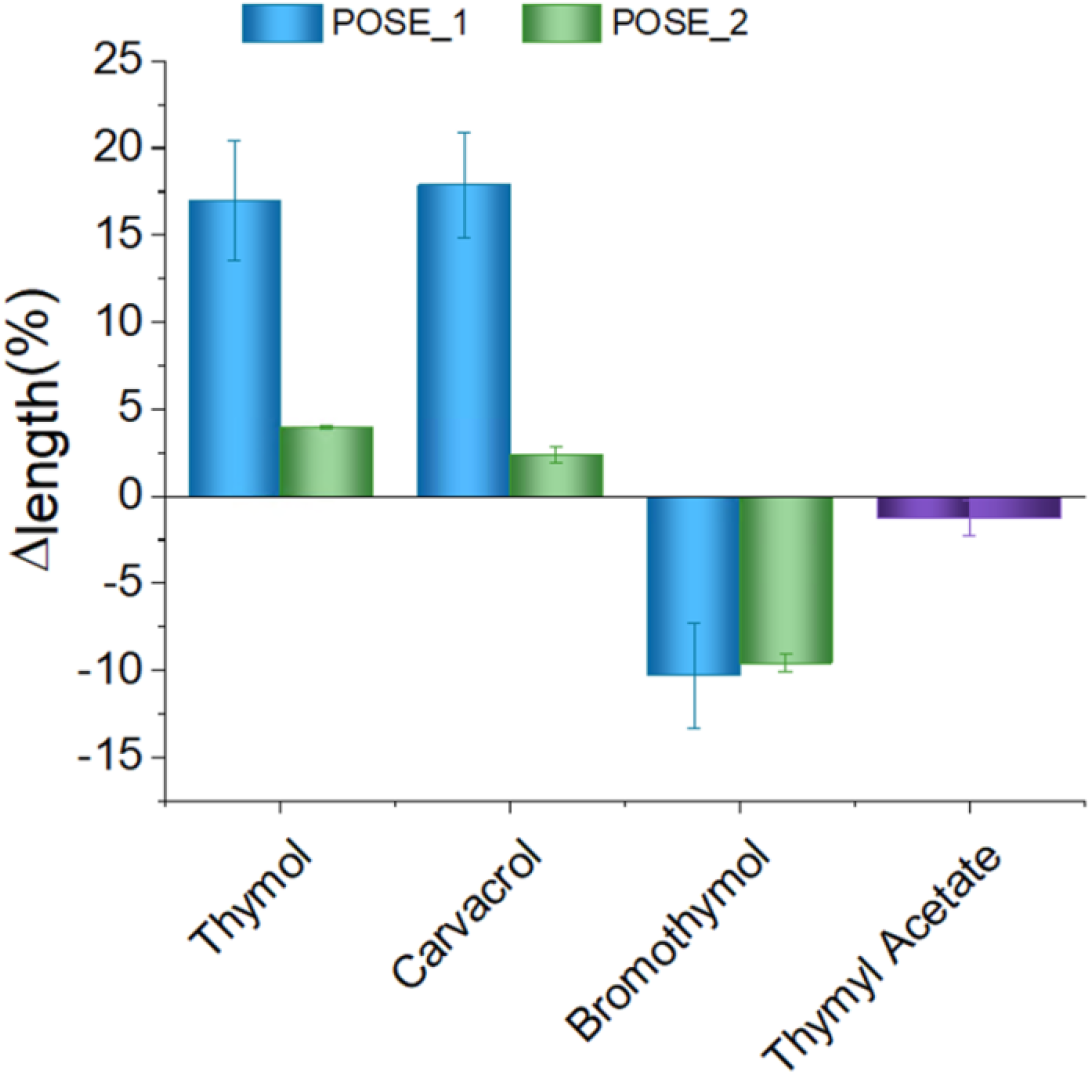
Changing in lipids PO4-—N+ percentage of variation in presence of each compound, setting the POPC one as reference (0 in the scale).

Thymol and carvacrol POSE_1 increase the PO4-—N+ distance of 14%, while POSE_2 had slightly or negligible effect (Figure 6, and Table S2). Contrary, bromothymol POSE_1 and POSE_2 induced a remarkable shrinkage of PO4-—N+ distance, probably due to the presence of the strong electronegative bromine. Lacking any interaction, thymyl acetate did not significatively influence the PO4-—N+ dipole. Overall, the effect of bromothymol on the PO4-—N+ distance is significantly higher than that observed for the other molecules. This evidence suggests the possibility that, in addition to the higher affinity discussed above, bromothymol presence in the bilayer seems to remarkably affect the polar head dynamics.

#### 2.2.4 Potential reactivity and stability studies

Differences in compound reactivity were investigated through a QM approach, exploiting electrostatic potential map (EMP) and atomic potential tensor (APT) studies, Figure 7.

**Figure 7.**
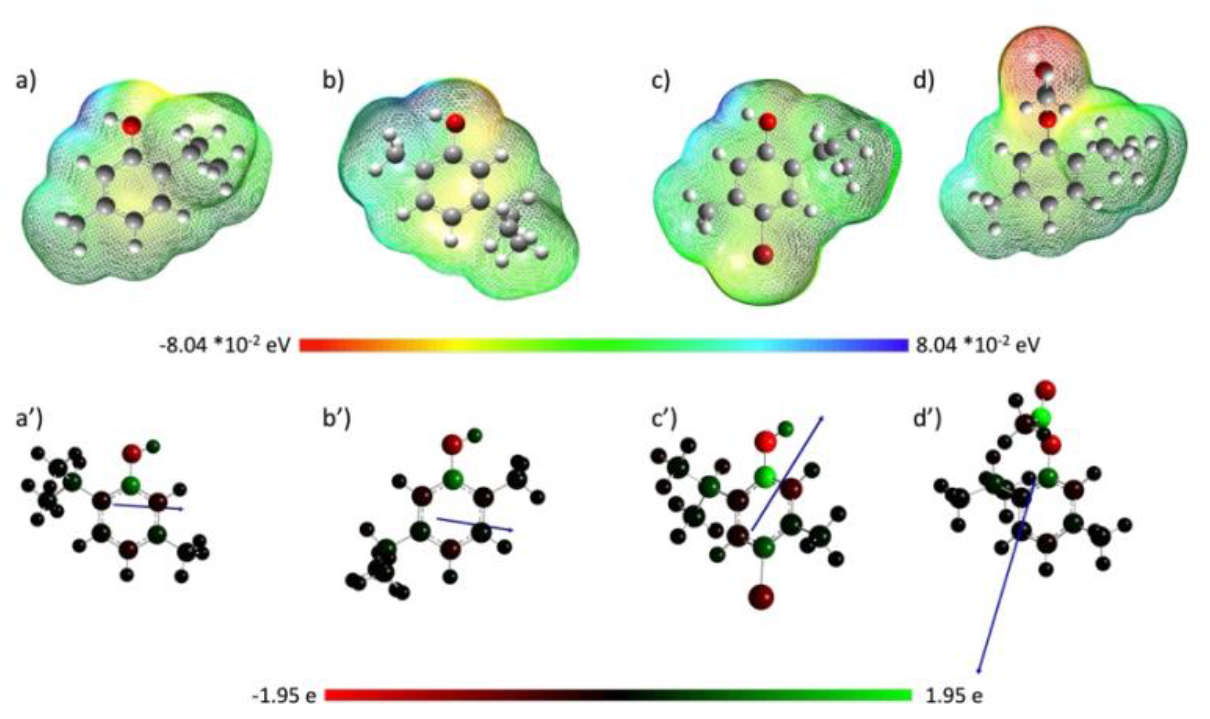
Surface electrostatic potential map (EPM) and atomic potential tensor (APT) of a-a’) Thymol; b-b’) Carvacrol; c-c’) Bromothymol; d-d’) Thymyl-acetate, in SMD solvation model. Geometry optimization performed with B3LYP/SMD/6-311 G+ (d,p).

EPM is a very useful tool in understanding a panel of molecular properties, sweeping from nucleophilicity, to surface local ionization energy. Moreover, it is suitable in investigating similarity between molecules behaviour [45]. Results corroborate the possibility that aromatic ring established π−π or π−σ interactions, while -OH group, when present, acted as acceptor and/or donor in hydrogen bond interactions. EMP validate the possibility of thymyl acetate, (Figure 7, panel d), in acting only as H-bond acceptor.

Recently, atomic charge analysis demonstrated its reliability in better determining the most probable site to undergo a nucleophilic attack [46]. Thus, the analysis of the atomic charges emphasizes differences in chemical and structural characteristic, predicting the intermolecular and intramolecular bonding, and reverting a more detailed picture of molecules behaviour [47]. Thus, atomic potential tensor (APT) was investigated for each compound, Figure 7, panel a’-d’) and table S3. Contrary to other approaches, APT produce physical interpretation of the charge localization, due to its correlation at the first derivate of cartesian coordinate of dipolar momentum. It showed that the hydrogen belonging to bromothymol has a more acid behaviour, accentuating a major chance in established hydrogen bond with respect to thymol and carvacrol. Furthermore, as showed in table S3, the bromothymol charge density inversion, in carbon C3 and C4, allows a stronger interaction with the nucleophilic and/or electrophilic moieties of the lipid counterpart. Overall, these data are coherent with the stronger activity with respect to that of thymol and carvacrol. Eventually, compound stability and potential reactions with different substrates were also evaluated through HOMO/LUMO orbitals analysis [21], in vacuum, keeping constant the functional and the basis. Table S4 values suggested that bromothymol, having a lower HOMO-LUMO gap (ΔE=-0.21), showed a higher reactivity and a lower kinetic stability with respect to the other compounds [22]. This was also reinforced by FMO map analysis (Figure S5, panel c), where the presence of the electron withdrawing substituent lead to a delocalization of the HOMO between the aromatic ring and the halogen atom. Furthermore, the FMO well agreed with the EPM outputs in term of interaction/reactivity of the aromatic ring, and -OH group, as well as in predicting possible changes in geometry hindrance of isomers.

## 3. Materials and Methods

### 3.1 In vivo toxicity determination

The antimycotic activity of thymol and bromothymol was determined evaluating the Minimum Inhibitory Concentration (MIC) on eight different strains of mould and yeast. They were previously isolated and identified based on macro and microscopic characteristics and properly preserved. Antimycotic activity was determined according to the serial dilution method. Briefly, thymol and bromothymol were dissolved in dimethylsulphoxide (DMSO). For each titled concentrations, different volumes were added to 20 ml of Sabouraud Dextrose Agar SDA (Liofilchem®, Italia). Thus, serial dilutions were obtained: 3.5, 7, 14, 28, 56, 112, 225, 450, 900 μg/ml. Each concentration was linked to a Perti’s dishes with an equal amount of DMSO, as control. Each mycotic species was inoculated in humidity control environment (37%) and at 27°C for 7 days.

### 3.2 Computational quantum mechanism method

DFT calculations were performed using Gaussian 16rev. A. 03 [31]. Because the electronic pattern and its distribution affect the biological activity of drug-like molecules, geometry and frequency optimization have been performed in the vacuum and in continuum solvent. The true minima of each species were selected avoiding negative frequency. Calculations were computed using B3LYP/SMD/6-311G+(d,p). B3LYP was selected for its reliability in this kind of application [48], while 6-311 G+(d,p) was selected as basis set, for its reliability in modelling hydrogen bond interaction [48]. SMD was selected as continuum solvent model to mimic water molecules interactions with compounds, proving its capability in modelling appropriately the solvation of charged and uncharged solutes. After geometry optimization, Hessian analysis indicate the absence of imaginary frequencies. Electrostatic potential maps (EPM) and frontiers molecular orbital (HOMO and LUMO) were computed with Gaussian 16 rev. and visualized trough Gauss-View 6.0.16. HOMO was pointed as the region where compounds could donate electrons during the interaction with the solvent and/or the lipids moiety. While the capacity of those compounds to accept electrons from the partner solvent and/or lipids is signified by LUMO energy. Partition coefficient was evaluated according to the literature [48] using the same level of theory exploited in MPE and FMO determination (B3LYP/SMD/6-311G+(d,p)). After preliminary computations in vacuum, geometry and frequency optimization were conducted in water and heptane, respectively. LogP was calculated according to equation 1. Single point energy was calculated with CAM-B3LYP, in vacuum, and 6-31 as basis set, activating two fragmentation groups option. Final structure of the dynamics output was extracted by pdb file format, and its backbone was re-draw in Gauss-View, to generate the input file.

### 3.3 Molecular dynamics simulation

All simulations were performed using GROMACS (version 5.1.1). For each compound a set of ten independent boxes was build. Cube dimensions were fixed at 9 x 9 x 9 nm. 1 or 0 (for control) target, 128 POPC [23], 23 Na+ Cl-ion pair, and 7500 water (SPCmodel [23]) molecules were randomly inserted. Molecules parameters were generated using ATB [49]. According to those conditions, a Gromos54a7_atb was applied to minimize the system energy. Thereafter, the MBA protocol was adopted with a few small variations [23]. Briefly, the temperature was fixed at 310°K, avoiding lipids jellification. Temperature and pressure were controlled through a Berendsen coupling. For the pressure, anisotropic conditions were used. This state was kept constant for 100 ns with a time step equal to 0.002 fs. Thereafter, each box was orientated with the axis perpendicular to the membrane plane parallel to the z axes and a second 100 ns long dynamics was carried out. In this second step, a semi-isotropical condition was applied using a Parinello−Rahman barostat. Only defect-free dynamics were exploited for the analysis.

### 3.4 Data Analysis

Density, RDF, Hydrogen bond formation, pair-distance and order parameter (SCD), were calculated using GROMACS tools (GROMACS) while lipid number for leaflet, and 2D lateral membrane thickness were calculated using the membrane plug-in implemented in VMD [50]. Bilayer thickness was expressed as the distance between the peaks of the lipid phosphorus atoms (P8 type in our system). Molecules visualization into the double layer were performed using free Maestro version. The standard error of the mean was obtained from seven independent MD simulations. Graphs and interpolations were produced with Origin [51].

## 4. Conclusions

The compelling needs in developing new antimycotic drugs has raised interest in natural compounds [5]. In this work, thymol antifungal in vivo efficacy was compared to the one of its brominated analogues. Bromothymol proved to be up to six-time more effective than the natural counterpart, even on difficult-to-treat fungi, showing an overall average MIC approximately equal to 42 μg/ml. To better identify similarity and differences in the mechanism of action, a detailed in silico characterization on four, natural and tailored, monoterpenoids was performed. A deep investigation was afforded on the well-known thymol, carvacrol and thymyl acetate, in parallel with the less studied bromothymol, to corroborate output. LogP_Hept/water_ suggested that bromothymol was the most hydrophobic compound, letting hypothesize its penetration into the bilayer. MBA investigation, first performed on this type of compounds, exhibited remarkable accordance with other previous results [9, 14, 16,42], and helped in clarifying the overall mechanism of action. Apparently, only slightly differences among molecules behaviour were noticeable. Indeed, all compounds bearing the -OH decreased membrane thickness and increased lipids order parameter, possibly acting on the membrane diffusion and permeability, as observed experimentally [17, 44]. The two possible poses of thymol and carvacrol, reported in different papers [14-16], were predicted also in herein. Referring to thymol and carvacrol, only POSE_1, the most energetically favoured in the former, disturbed the relative length of 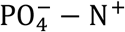, increasing the distance between the dipole partners (Δl=1Å, table S2). Contrary, bromothymol, independently from the placement, aroused a shrinkage of the lipid head (Δl=-1Å, table S2), probably because of the strong electronegative bromine, suggesting a stronger perturbation of the polar heads dynamics with respect to the other three analysed compounds. QM analysis of EPM, APT and FMO also reinforce this evidence, emphasized the differences in nucleophilicity of the -OH group, the charges distribution on the aromatic core, and the reactivity of the compounds. Overall, the reported computational findings were in excellent agreement with the experimental results [42], and they clarified, for the first time, the mechanisms of action of thymol, carvacrol, and bromothymol, explaining their different efficiency in antimicrobial activity. Thus, the polar distortion of the lipid head triggers a “domino” effect which influences the whole membrane stability, as also demonstrated for other compounds [43].

## Supplementary Materials

The following supporting information can be downloaded at:

## References

[1] Meis, J. F., Chowdhary, A., Rhodes, J. L., Fisher, M. C. , Verweij, P. E. Philos Trans R Soc Lond B Biol Sci. 2016, 371, 20150460.

[2] Kan J, Liu T, Ma N, Li H, Li X, Wang J, Zhang B, Chang Y, Lin J. PLoS ONE. 2017, 12:e0184988. doi: 10.1371/journal.pone.0184988.

[3] Sudharshana TN, Venkatesh HN, Nayana B, Manjunath K, Mohana DC. Mycology 2018, 10, 40–48.

[4] Piombino, C., Lange, H., Sabuzi, F., Galloni, P., Conte, V., Crestini, C. Molecules 2020, 25, 866

[5] Floris, B.; Galloni, P.; Conte, V.; Sabuzi, F. Biomolecules 2021, 11, 1325.

[6] Mangena, T.; Muyima, N. Y. O. Lett. Appl. Microbiol. 1999, 28 (4), 291–296.

[7] Skandamis, P.; Tsigarida, E.; Nychas, G. E. Food Microbiol. 2002, 19 (1), 97–103.

[8] Gutierrez, J.; Barry-Ryan, C.; Bourke, P. Food Microbiol. 2009, 26 (2), 142–150.

[9] Kowalczyk, A., Przychodna, M., Sopata, S., Bodalska, A., Fecka, I. Molecules 2020, 25, 4125.

[10] Yuan, Z.; Dai, Y. P Ouyang, P.; Rehman, T.; Hussain, S.; Zhang, T.; Yin, Z.; Fu, H., Lin, J.; He, C., Lv, C.; Liang, X.; Shu, G.; Song, X.; Li, L.; Zou Y., Yin, L. Microorganisms 2020, 8, 99

[11] Tohidpour, A.; Sattari, M.; Omidbaigi, R.; Yadegar, A.; Nazemi, J. Phytomedicine, 2010, 17, 2, 142–145.

[12] Neri, F.; Mari, M.; Brigati, S. Plant Pathol 2006, 55, 100–105.

[13] Coppo, E.; Marchese, A. Current Pharmaceutical Biotechnology, 2014, 15, 4, 380–390.

[14] Miguel, V.; Villarreal, M.A.; Garcia, D.A. PLOS ONE 2016, 14(6): e0218042

[15] Reiner, G. N.; Perillo, M.A; Garcia, D.A. Colloids Surf B Biointerfaces. 2013; 101, 61–7. Epub 2012/07/17

[16] Ferreira, J. V. N., Capello, T. M., Siqueira, L. J. A., Lago, J. H. G., Caseli, L. Langmuir 2016, 32, 3234–3241

[17] Zhang, J., Ma, S., Du, S., Chen, S., Sun, H. J Food Sci Technol 2019, 56(5), 2611–2620.

[18] Lazarević, J., Marković, A., Šmelcerović, A., Stojanović, G., Ciuffreda, P., Santaniello, E. Acta Chim. Slov. 2022, 69, 571–583

[19] S. Pezzola, M. Venanzi, P. Galloni, V. Conte, F. Sabuzi. Chemistry, 2024, X, YYY

[20] S. Pezzola, S. Tarallo, A. Iannini, M. Venanzi, P. Galloni, V. Conte, F. Sabuzi, Molecules 2022, 27, 8590

[21] Zhan, C.G., Nichols, J. A., Dixon, D. A. J. Phys. Chem. A, 2003, 107, 4184–4195

[22] Miar, M., Shiroudihttps, A., Pourshamsian, K., Oliaey, A. R., Hatamjafari, F. Journal of Chemical Research 2021, 45, 147–158

[23] Farrotti, A., Bocchinfuso, G., Palleschi, A., Rosato, N., Salnikov, E.S., Voievoda, N., Bechinger, B., Stella, L. Biochimica et Biophysica Acta 2015, 1848, 581–592

[24] Bobone, S., Bocchinfuso, G., Park, Y., Palleschi, A., Hahm, K.-S., Stella, L. J. Pept. Sci. 2013, 19, 758–769.

[25] Esteban-Martín, S. Salgado, J. Biophys. J. 2007, 92, 903–912.

[26] Orioni, B., Bocchinfuso, G., Kim, J.Y., Palleschi, A., Grande, G., Bobone, S., Park, Y., Kim, J.I., Hahm, K.-S., Stella, L. Biochim. Biophys. Acta 2009, 1788, 1523–1533

[27] Wang, K., Yan, J., Liu, X., Zhang, J., Chen, R., Zhang, B., Dang, W., Zhang, W., Kai, M., Song, J., Wang, R. Toxicology, 2011, 288, 27–33

[28] Fowler, P. W., Hélie, J., Duncan, A., Chavent, M., Koldsø, H., Sansom, M. S. P. Soft Matter, 2016, 12, 7792.

[29] Bobone, S., Gerelli, Y. De Zotti, M., Bocchinfuso, G., Farrotti, A., Orioni, B., Sebastiani, F., Latter, E., Penfold, J., Senesi, S., Formaggio, F., Palleschi, A., Toniolo, C., Fragneto, G., Stella, L. Biochimica et Biophysica Acta, 2013, 1828, 1013–1024

[30] Barman, S., Mukherjee, S., Jolly, L., Troiano, C., Grottesi, A., Basak, D., Calligari, P., Bhattacharjee, B., Bocchinfuso, G.,Stella, L. Haldar, J. Chem. Sci., 2023, 14, 4845

[31] Uygun, B. A., Bou-Akl, T., Albanna, M., Matthew, H. W. T. Acta Biomater. 2010, 6, 2126–2131.

[32] Wang, K., SJiang, S., Pu, T., Fan, L., Su, F., Ye, M. Natural Product Research 2018 DOI: 10.1080/14786419.2017.1419232

[33] Wang, K., SJiang, S., Pu, T., Fan, L., Su, F., Ye, M. Natural Product Research 2018, DOI: 10.1080/14786419.2018.1480618

[34] Sabuzi, F. Churakova, E., Galloni, P., Wever, R., Hollmann, F., Floris, B., Conte, V. Eur. J. Inorg. Chem. 2015, X, 3519–3525

[35] Avdeef, A., K. J. Box, J. E. Comer, C. Hibbert, and K. Y. Tam. Pharmaceutical Research, 1998, 15 (2): 209–15. doi:10.1023/a:1011954332221.

[36] M. Haritha, C. H. Suresh. J Comput Chem. 2022, 43, 477–490.

[37] Mayer, Peter T., and Bradley D. Anderson. 2002. Journal of Pharmaceutical Sciences 91 (3): 640–46. doi:10.1002/jps.10067.

[38] Bannan, C. C., Calabró, G., Kyu, D. Y., Mobley, D. L. J. Chem. Theory Comput. 2016, 12, 4015–4024

[39] PubChem. Available online: https://pubchem.ncbi.nlm.nih.gov (accessed on 10 March 2024)

[40] Hawash, M. , Jaradat, N., Abualhasan, M., Süküroglu, M. K., Qaoud, M. T., Kahraman, D. C., Daraghmeh, H., Maslamani, L., Sawafta, M., Ratrout, A., Issa, L. BMC Chemistry 2023, 17, 11

[41] S. Pezzola, S. Tarallo, A. Iannini, M. Venanzi, P. Galloni, V. Conte, F. Sabuzi, Molecules 2022, 27, 8590

[42] Y. Lyu, N. Xiang, J. Mondal, X. Zhu, G. Narsimhan, J Phys Chem B 2018, 122, 2341–2354.

[43] U. Schote, J. Seelig, Biochimica et Biophysica Acta, 1998, 1415, 135–146

[44] B. J. Zwolinski, H. Eyring, C. E. Reese, J. Phys. Chem. 1949, 53, 9, 1426–1453

[45] P.C Mishra, A. Kumar, Theoretical and Computational Chemistry, Vol 3, Elsevier Science B.V., 1996.

[46] Rangel-Galván, M., Castro, M. E., Perez-Aguilar, J. M., Caballero, N. A., Rangel-Huerta, A., Melendez, F. J. Molecules 2022, 27, 414.

[47] North SC, Jorgensen KR, Pricetolstoy J and Wilson AK. Front. Chem., 2023, 11:1152500.

[48] M. A. Nedyalkova, S. Madurga, M. Tobiszewski, V. Simeonov. J. Chem. Inf. Model. 2019, 59, 2257–2263

[49] Stroet M, Caron B, Visscher K, Geerke D, Malde AK, Mark AE. J Chem. Theory and Comput., 2018, 14, 5834–5845.

[50] Guixà-González, R.; Rodriguez-Espigares, I.; Ramírez-Anguita, J. M.; Carrió-Gaspar, P.; Martinez-Seara, H.; Giorgino, T.; Selent, J. Bioinformatics, 2014, 30, 1478–1480.

[51] Origin(Pro), Version 2021b. OriginLab Corporation, Northampton, MA, USA

