## supplementary info for "New insight in cyclic monoterpenoids mechanism of action: an in silico approach"

### Supporting Information

#### Contents


Figure 1 S2

Figure 2 S3

Figure 3 S4

Optimized cartesian coordinates (in Angstroms) S5


**Table S3.** MIC value (mg/ml) for each compound on selected yeast and mould species.

| Specie | Thymol (mg/ml) | Bromothymol (mg/ml) |
|---|---|---|
| *Alternaria alternata* | 225 ±0 | 42±14 |
| *Aspergillus fumigatus* | 225 ±0 | 42±14 |
| *Aspergillus niger* | 168.5 ±56.5 | 28±0 |
| *Fusarium oxysporum* | 168.5 ± 56.8 | 56±0 |
| *Microsporum canis* | 56 ±0 | 28±0 |
| *Penicillium glaucum* | 225 ±0 | 56±0 |
| *Penicillium chrysogenum* | 168.5 ± 56.5 | 56±0 |
| *Poeciliomyces liliacinus* | 84 ±28 | 14±0 |
| *Candida albicans* | 112 ±0 | 42±14 |
| *Rhodotorula mucilaginosa* | 112 ±0 | 42±14 |
| *Malassezia pachydermatis* | 56 ±0 | 14±±0 |

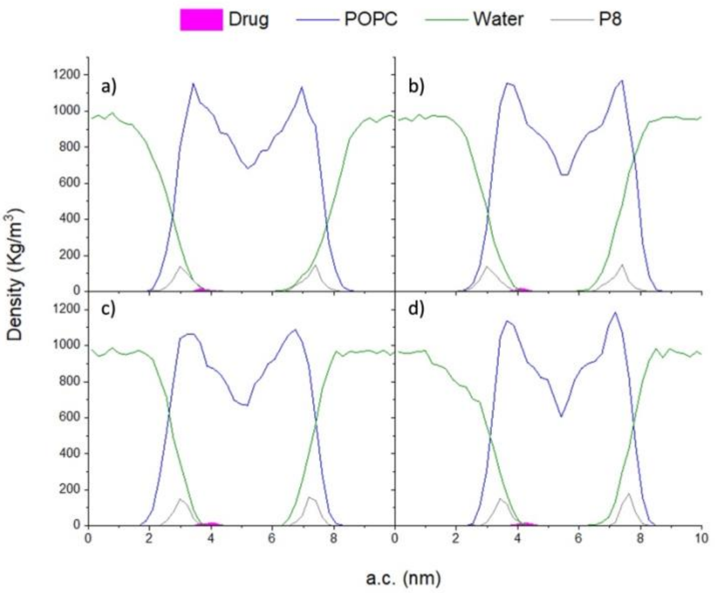


**Figure S1.** Compounds final position into the bilayer. Panels represented the average of seven independent dynamics, after 100 ns of semi-isotropic dynamics. Panel: a). thymol; b) carvacrol; c) bromothymol; d) thymyl acetate.

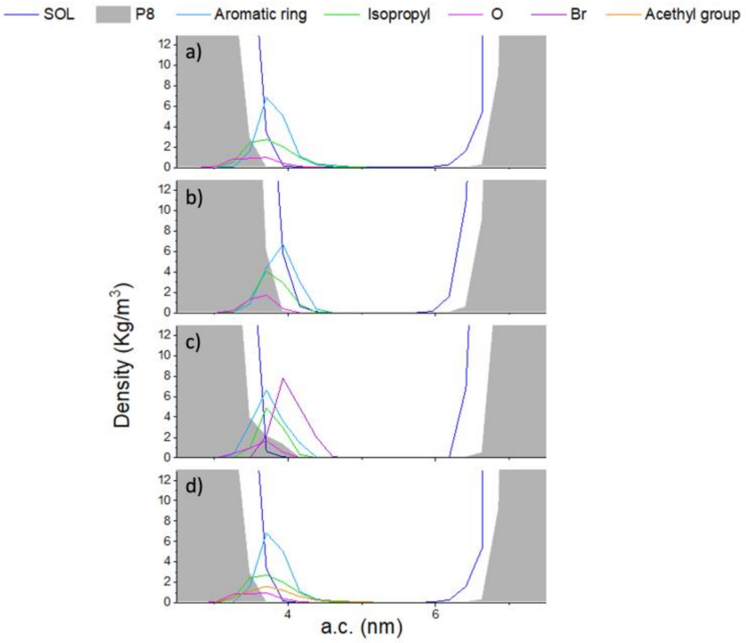


**Figure S2.** Average density of the three regions of interest identified in QM into the double layer. Panels: a) thymol; b) carvacrol; c) bromothymol; d) thymyl acetate. O: oxygen bond to the phenyl group.

**Scheme 1.** Schematic representation of the lipid groups considered in the analysis.

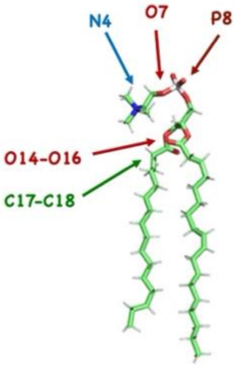


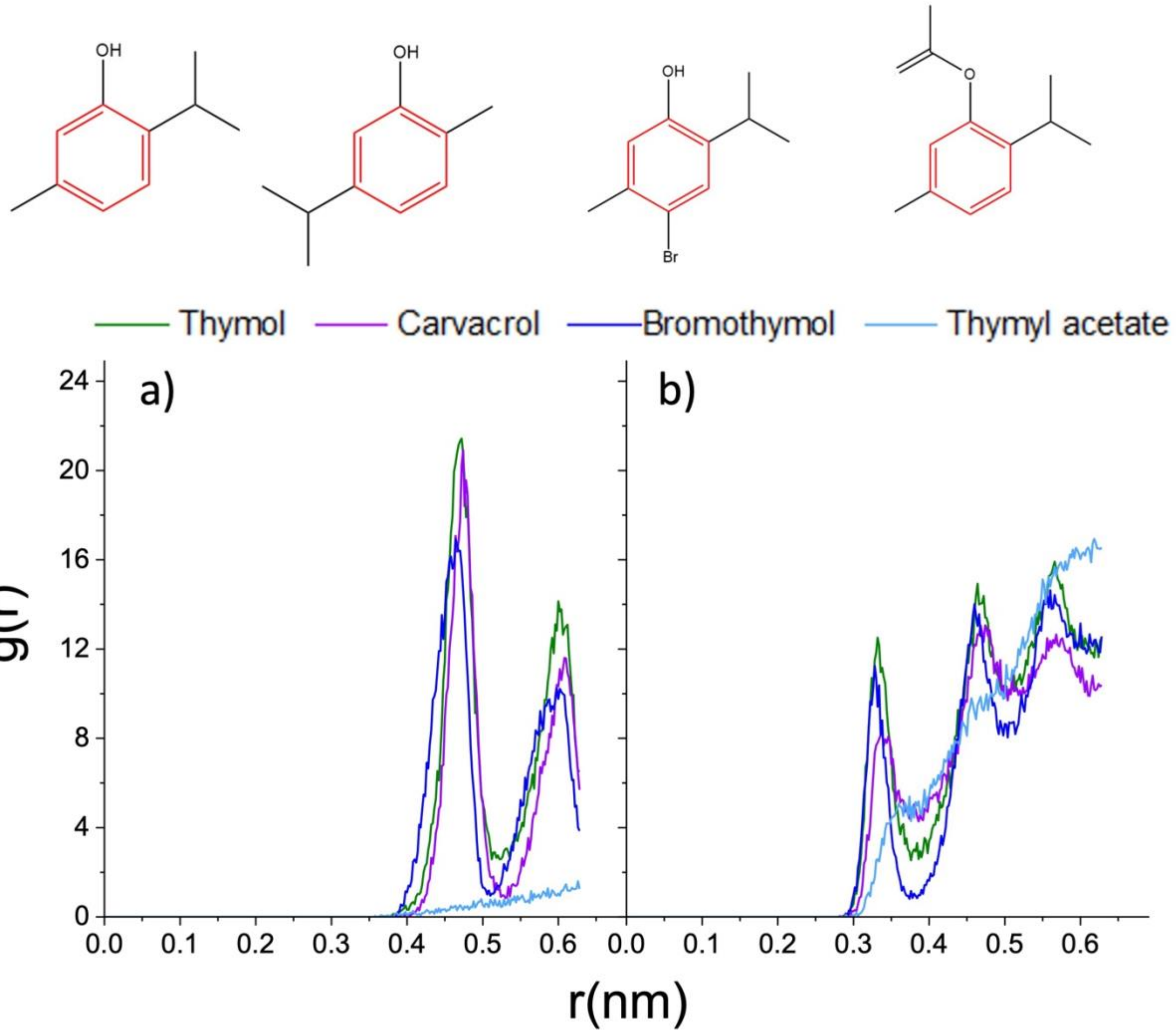


**Figure S3.** Sum of the radial distribution function g(r) of at least seven dynamics for each compound, with respect to the common feature, the aromatic ring. Panels: a) RDF of the aromatic ring with P8; b) RDF of the aromatic ring with O14-16, respectively.

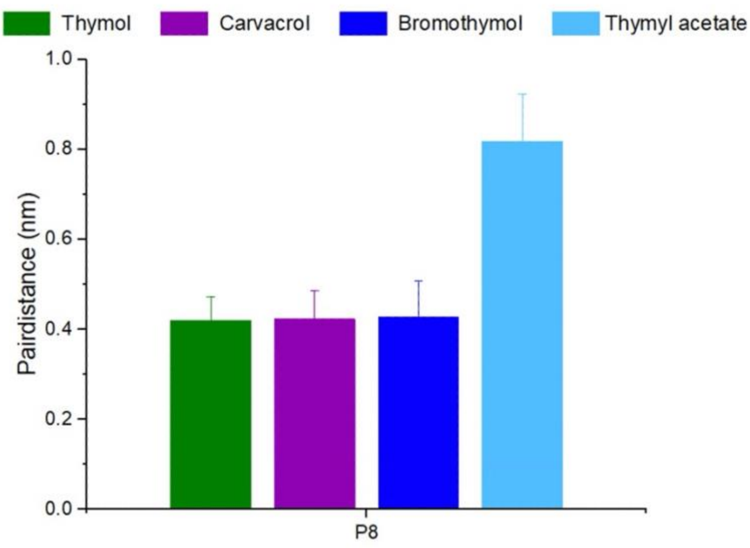


**Figure S4.** Average distance of the oxygen of monoterpenoids from P8, linkable to compounds penetration into the double layer.

**Table S2.** Average variation of the $PO_4^- - N^+$ relative distance in presence or not of compound.

| **Compound** | | $PO_4^- - N^+$ |
|---|---|---|
| POPC | | 4.97 ± 0.07 |
| Thymol | POSE_1 | 5.81 ± 0.40 |
| | POSE_2 | 5.17 ± 0.00 |
| Carvacrol | POSE_1 | 5.86 ± 0.90 |
| | POSE_2 | 5.09 ± 0.61 |
| Bromothymol | POSE_1 | 4.46 ± 0.22 |
| | POSE_2 | 4.49 ± 0.30 |

**Table S3.** Calculation of the atom charges with APT. Geometry was optimized using B3LYP/SMD/6-311G+(d,p).

| Compounds | Atoms Charge *e* | | | | | | | | | | | |
|---|---|---|---|---|---|---|---|---|---|---|---|---|
| | -OH | | Aromatic ring | | | | | | Isopropyl | | | Methyl |
| | H1 | O2 | C1 | C2 | C3 | C4 | C5 | C6 | C7 | C8 | C9 | C10 |
| Thymol | 0.40 | -1.05 | 0.802 | -0.13 | 0.04 | -0.25 | 0.21 | -0.27 | 0.24 | 0.49 | 0.05 | 0.06 |
| Carvacrol | 0.42 | -1.07 | 0.806 | -0.24 | 0.15 | -025 | 0.05 | -0.01 | 0.24 | 0.05 | 0.05 | 0.06 |
| Bromothymol | 0.40 | -1.08 | 0.826 | -0.10 | -0.10 | 0.30 | 0.07 | -0.24 | 0.23 | 0.05 | 0.04 | 0.05 |
| Thymyl acetate | | -1.4 | 0.690 | -0.06 | -0.06 | -0.11 | 0.13 | -0.20 | 0.23 | 0.06 | 0.05 | 0.05 |

**Table S4.** Frontier orbital energies of the four compounds. LUMO: low unoccupied molecular orbital; HOMO: Highest occupied molecular orbital; ΔE: difference in orbital energy between LUMO and HOMO. Data were obtained by the geometry optimization using B3LYP/SMD/6-311G+(d,p).

| Compound | LUMO (eV) | HOMO (eV) | ΔE (eV) |
|---|---|---|---|
| Thymol | -0.01436 | -0.22560 | -0.21124 |
| Carvacrol | -0.01073 | -0.2257 | -0.21497 |
| Bromothymol | -0.01971 | -0.22477 | -0.20506 |
| Thymyl-acetate | -0.02052 | -0.24256 | -0.22204 |

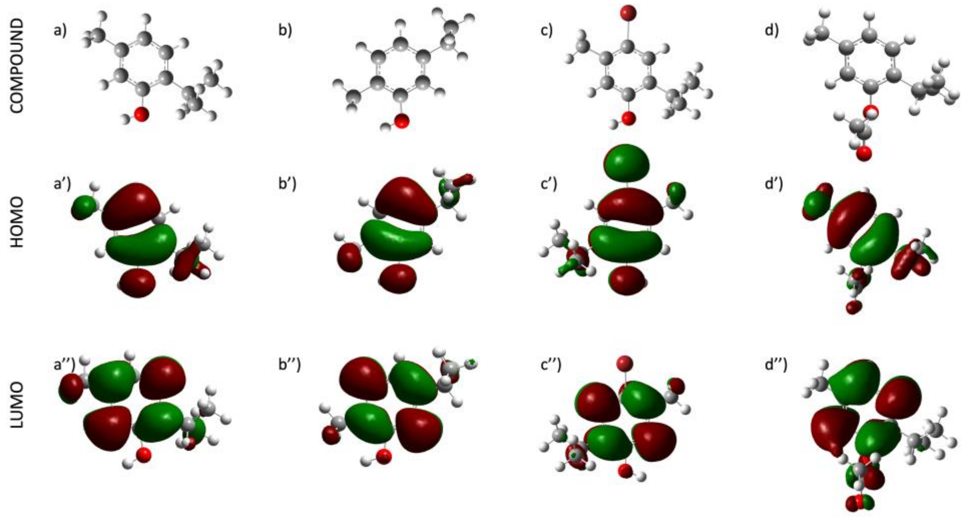


**Figure S5**. Frontier molecular orbital map of compounds: a), a'), and a'') thymol; b), b'), and b'') carvacrol; c), c'), and c'') bromothymol; d), d'), and d'') thymil-acetate. Geometry optimized with B3LYP/SMD/6-311G+ (d, p)

**Optimized cartesian coordinates (in Angstroms)**

**B3LYP 6-311G+(d,p) vacuum**

**Thymol**
Free Energy -464.683856
Stoichiometry C10H14O

Framework group  C1[X(C10H14O)]
Deg. of freedom    69
Full point group                 C1      NOp   1
Largest Abelian subgroup         C1      NOp   1
Largest concise Abelian subgroup C1      NOp   1
Standard orientation:

| Center Number | Atomic Number | Atomic Type | Coordinates (Angstroms) X | Y | Z |
|---|---|---|---|---|---|
| 1 | 6 | 0 | -1.487556 | -1.406607 | 0.114812 |
| 2 | 6 | 0 | -0.096818 | -1.353538 | 0.021544 |
| 3 | 6 | 0 | 0.587900 | -0.147104 | -0.134635 |
| 4 | 6 | 0 | -0.195760 | 1.016962 | -0.208257 |
| 5 | 6 | 0 | -1.580524 | 0.972318 | -0.095549 |
| 6 | 6 | 0 | -2.256898 | -0.243710 | 0.062391 |
| 7 | 1 | 0 | -1.973549 | -2.369104 | 0.235786 |
| 8 | 1 | 0 | 0.462150 | -2.281319 | 0.075478 |
| 9 | 1 | 0 | -2.123689 | 1.910979 | -0.129383 |
| 10 | 8 | 0 | 0.425267 | 2.262155 | -0.313936 |
| 11 | 1 | 0 | 0.466566 | 2.527689 | -1.238657 |
| 12 | 6 | 0 | 2.100309 | -0.054487 | -0.191404 |
| 13 | 1 | 0 | 2.332574 | 0.698412 | -0.955147 |
| 14 | 6 | 0 | 2.677745 | 0.461751 | 1.149619 |
| 15 | 1 | 0 | 3.748792 | 0.247475 | 1.217796 |
| 16 | 1 | 0 | 2.532293 | 1.538101 | 1.257445 |
| 17 | 1 | 0 | 2.185785 | -0.022977 | 1.998452 |
| 18 | 6 | 0 | 2.772392 | -1.364095 | -0.620897 |
| 19 | 1 | 0 | 2.364655 | -1.744605 | -1.562926 |
| 20 | 1 | 0 | 3.844142 | -1.197808 | -0.762769 |
| 21 | 1 | 0 | 2.664271 | -2.149136 | 0.135631 |
| 22 | 6 | 0 | -3.759620 | -0.280534 | 0.174324 |
| 23 | 1 | 0 | -4.115391 | 0.363101 | 0.985614 |
| 24 | 1 | 0 | -4.237719 | 0.069605 | -0.747295 |
| 25 | 1 | 0 | -4.120044 | -1.293392 | 0.369771 |

Rotational constants (GHZ):           2.0108567           0.7387148           0.5884603

**Carvacrol**
Free Energy -464.683159
Stoichiometry    C10H14O
Framework group  C1[X(C10H14O)]
Deg. of freedom    69
Full point group                 C1      NOp   1
Largest Abelian subgroup         C1      NOp   1
Largest concise Abelian subgroup C1      NOp   1
Standard orientation:

| Center Number | Atomic Number | Atomic Type | Coordinates (Angstroms) X | Y | Z |
|---|---|---|---|---|---|
| 1 | 6 | 0 | -0.407958 | -1.339176 | -0.041140 |
| 2 | 6 | 0 | 0.955176 | -1.598250 | 0.104528 |
| 3 | 6 | 0 | 1.871084 | -0.546611 | 0.084790 |
| 4 | 6 | 0 | 1.424079 | 0.764736 | -0.081867 |
| 5 | 6 | 0 | 0.061300 | 1.023625 | -0.227969 |
| 6 | 6 | 0 | -0.854845 | -0.028353 | -0.207215 |
| 7 | 1 | 0 | 1.307105 | -2.631714 | 0.236211 |
| 8 | 1 | 0 | -0.291306 | 2.057011 | -0.359339 |
| 9 | 8 | 0 | 2.363410 | 1.842764 | -0.102138 |
| 10 | 1 | 0 | 2.652735 | 1.998784 | -1.004108 |
| 11 | 6 | 0 | 3.375676 | -0.832386 | 0.246452 |
| 12 | 1 | 0 | 3.894715 | -0.520791 | -0.635821 |
| 13 | 6 | 0 | -2.359422 | 0.257844 | -0.368276 |
| 14 | 1 | 0 | -2.543725 | 0.659799 | -1.342629 |
| 15 | 1 | 0 | -1.110506 | -2.146073 | -0.025536 |
| 16 | 1 | 0 | 3.752314 | -0.293773 | 1.090810 |
| 17 | 1 | 0 | 3.525395 | -1.881153 | 0.396693 |
| 18 | 6 | 0 | -2.805781 | 1.274186 | 0.699157 |
| 19 | 1 | 0 | -2.486441 | 2.254706 | 0.413670 |
| 20 | 1 | 0 | -3.872266 | 1.257037 | 0.784090 |

| | | | | | |
|---|---|---|---|---|---|
| 21 | 1 | 0 | -2.368766 | 1.016975 | 1.641368 |
| 22 | 6 | 0 | -3.152381 | -1.050817 | -0.194428 |
| 23 | 1 | 0 | -4.199405 | -0.847442 | -0.279743 |
| 24 | 1 | 0 | -2.861442 | -1.747542 | -0.952603 |
| 25 | 1 | 0 | -2.947248 | -1.466732 | 0.769852 |

---------------------------------------------------------------------
Rotational constants (GHZ): 2.2015963 0.6875189 0.5467780

**-Bromothymol**
Free Energy -3038.237361
Stoichiometry C10H13BrO
Framework group C1[X(C10H13BrO)]
Deg. of freedom 69
Full point group C1 NOp 1
Largest Abelian subgroup C1 NOp 1
Largest concise Abelian subgroup C1 NOp 1
Standard orientation:

| Center Number | Atomic Number | Atomic Type | Coordinates (Angstroms) X | Y | Z |
|---|---|---|---|---|---|
| 1 | 6 | 0 | -0.933837 | 0.023545 | -0.032152 |
| 2 | 6 | 0 | 0.237760 | -0.726714 | -0.136764 |
| 3 | 6 | 0 | 1.474863 | -0.084756 | -0.188630 |
| 4 | 6 | 0 | 1.540674 | 1.308164 | -0.137077 |
| 5 | 6 | 0 | 0.369286 | 2.058197 | -0.032968 |
| 6 | 6 | 0 | -0.868052 | 1.415849 | 0.019896 |
| 7 | 1 | 0 | 0.185607 | -1.824395 | -0.177000 |
| 8 | 1 | 0 | 0.420821 | 3.155997 | 0.007675 |
| 9 | 8 | 0 | 2.809221 | 1.966079 | -0.190353 |
| 10 | 1 | 0 | 3.036080 | 2.151149 | -1.104620 |
| 11 | 6 | 0 | 2.768206 | -0.912846 | -0.303216 |
| 12 | 1 | 0 | 3.267182 | -0.670219 | -1.218124 |
| 13 | 6 | 0 | 3.691502 | -0.590049 | 0.886291 |
| 14 | 1 | 0 | 4.176813 | -1.485458 | 1.214340 |
| 15 | 1 | 0 | 4.428112 | 0.124103 | 0.582493 |
| 16 | 1 | 0 | 3.111092 | -0.184510 | 1.688514 |
| 17 | 6 | 0 | 2.420102 | -2.412933 | -0.290527 |
| 18 | 1 | 0 | 1.784085 | -2.638522 | -1.120884 |
| 19 | 1 | 0 | 3.319261 | -2.988360 | -0.363310 |
| 20 | 1 | 0 | 1.915095 | -2.654186 | 0.621430 |
| 21 | 6 | 0 | -2.161157 | 2.244223 | 0.135116 |
| 22 | 1 | 0 | -2.099131 | 2.885262 | 0.989588 |
| 23 | 1 | 0 | -2.284888 | 2.836858 | -0.747139 |
| 24 | 1 | 0 | -2.997909 | 1.586106 | 0.242956 |
| 25 | 35 | 0 | -2.627774 | -0.855958 | 0.039631 |

---------------------------------------------------------------------
Rotational constants (GHZ): 1.1257593 0.4029779 0.3054390

**Thymyl Acetate**
Free Energy -617.342007
Stoichiometry C12H16O2
Framework group C1[X(C12H16O2)]
Deg. of freedom 84
Full point group C1 NOp 1
Largest Abelian subgroup C1 NOp 1
Largest concise Abelian subgroup C1 NOp 1
Standard orientation:

| Center Number | Atomic Number | Atomic Type | Coordinates (Angstroms) X | Y | Z |
|---|---|---|---|---|---|
| 1 | 6 | 0 | -0.203660 | 2.534386 | -0.088407 |
| 2 | 6 | 0 | -1.298286 | 1.683631 | -0.244879 |
| 3 | 6 | 0 | -1.130019 | 0.303338 | -0.136715 |
| 4 | 6 | 0 | 0.133566 | -0.226845 | 0.126864 |
| 5 | 6 | 0 | 1.227970 | 0.623741 | 0.282816 |
| 6 | 6 | 0 | 1.059301 | 2.004486 | 0.175567 |
| 7 | 1 | 0 | -0.336620 | 3.622624 | -0.173281 |
| 8 | 1 | 0 | -2.293991 | 2.101749 | -0.452220 |
| 9 | 1 | 0 | 2.223954 | 0.206200 | 0.490546 |
| 10 | 6 | 0 | -2.338380 | -0.635778 | -0.308542 |

| | | | | | |
|---|---|---|---|---|---|
| 11 | 1 | 0 | -2.175478 | -1.276294 | -1.150030 |
| 12 | 6 | 0 | -2.509257 | -1.490042 | 0.961353 |
| 13 | 1 | 0 | -3.551152 | -1.606480 | 1.175356 |
| 14 | 1 | 0 | -2.067328 | -2.452082 | 0.806179 |
| 15 | 1 | 0 | -2.028018 | -1.005112 | 1.784852 |
| 16 | 6 | 0 | -3.610319 | 0.201235 | -0.539151 |
| 17 | 1 | 0 | -3.494256 | 0.789702 | -1.425230 |
| 18 | 1 | 0 | -4.450986 | -0.451065 | -0.651774 |
| 19 | 1 | 0 | -3.769465 | 0.846628 | 0.299324 |
| 20 | 6 | 0 | 2.267764 | 2.943346 | 0.348075 |
| 21 | 1 | 0 | 2.735774 | 2.752683 | 1.291215 |
| 22 | 1 | 0 | 2.970030 | 2.769299 | -0.440234 |
| 23 | 1 | 0 | 1.937134 | 3.960382 | 0.313102 |
| 24 | 8 | 0 | 0.305787 | -1.642109 | 0.237632 |
| 25 | 6 | 0 | 1.641228 | -1.991164 | -0.136075 |
| 26 | 8 | 0 | 2.019451 | -3.190436 | -0.088484 |
| 27 | 6 | 0 | 2.619894 | -0.895670 | -0.598358 |
| 28 | 1 | 0 | 2.105696 | -0.201220 | -1.229419 |
| 29 | 1 | 0 | 3.008750 | -0.381335 | 0.255546 |
| 30 | 1 | 0 | 3.425220 | -1.343301 | -1.142396 |

---------------------------------------------------------------------

Rotational constants (GHZ): 0.7618127 0.6468214 0.3702056

- **B3LYP/SMD/6-311G+(d,p) Water**

**Thymol**

Free Energy -464.689543
Stoichiometry C10H14O
Framework group C1[X(C10H14O)]
Deg. of freedom 69
Full point group C1 NOp 1
Largest Abelian subgroup C1 NOp 1
Largest concise Abelian subgroup C1 NOp 1
Standard orientation:

| Center Number | Atomic Number | Atomic Type | Coordinates (Angstroms) X | Y | Z |
|---|---|---|---|---|---|
| 1 | 6 | 0 | -1.459966 | -1.420742 | 0.141815 |
| 2 | 6 | 0 | -0.068546 | -1.336043 | 0.052487 |
| 3 | 6 | 0 | 0.595199 | -0.118488 | -0.117770 |
| 4 | 6 | 0 | -0.212516 | 1.031069 | -0.193162 |
| 5 | 6 | 0 | -1.601528 | 0.957422 | -0.109733 |
| 6 | 6 | 0 | -2.249036 | -0.271354 | 0.059952 |
| 7 | 1 | 0 | -1.931509 | -2.388892 | 0.273426 |
| 8 | 1 | 0 | 0.511815 | -2.249192 | 0.115642 |
| 9 | 1 | 0 | -2.181233 | 1.874282 | -0.177172 |
| 10 | 8 | 0 | 0.418977 | 2.251773 | -0.354708 |
| 11 | 1 | 0 | -0.246494 | 2.949968 | -0.418524 |
| 12 | 6 | 0 | 2.112637 | -0.004859 | -0.178910 |
| 13 | 1 | 0 | 2.346108 | 0.836890 | -0.837616 |
| 14 | 6 | 0 | 2.690286 | 0.326148 | 1.211692 |
| 15 | 1 | 0 | 3.772689 | 0.477490 | 1.152161 |
| 16 | 1 | 0 | 2.244771 | 1.233946 | 1.626593 |
| 17 | 1 | 0 | 2.501107 | -0.494384 | 1.911492 |
| 18 | 6 | 0 | 2.800110 | -1.245892 | -0.763420 |
| 19 | 1 | 0 | 2.396462 | -1.504940 | -1.746449 |
| 20 | 1 | 0 | 3.870763 | -1.054248 | -0.879033 |
| 21 | 1 | 0 | 2.693069 | -2.117959 | -0.111817 |
| 22 | 6 | 0 | -3.753997 | -0.337870 | 0.139625 |
| 23 | 1 | 0 | -4.140564 | 0.329105 | 0.916078 |
| 24 | 1 | 0 | -4.213398 | -0.030645 | -0.805599 |
| 25 | 1 | 0 | -4.091257 | -1.351953 | 0.363027 |

---------------------------------------------------------------------

Rotational constants (GHZ): 2.0140033 0.7312023 0.5895998

**Carvacrol**

Free Energy -464.690462
Stoichiometry C10H14O
Framework group C1[X(C10H14O)]
Deg. of freedom 69
Full point group C1 NOp 1

Largest Abelian subgroup C1 NOp 1
Largest concise Abelian subgroup C1 NOp 1
Standard orientation:

| Center Number | Atomic Number | Atomic Type | Coordinates (Angstroms) X | Y | Z |
|---|---|---|---|---|---|
| 1 | 6 | 0 | -0.428545 | -1.241613 | 0.000001 |
| 2 | 6 | 0 | 0.931406 | -1.543961 | 0.000004 |
| 3 | 6 | 0 | 1.915535 | -0.548652 | 0.000002 |
| 4 | 6 | 0 | 1.470003 | 0.780392 | -0.000003 |
| 5 | 6 | 0 | 0.110926 | 1.095201 | -0.000006 |
| 6 | 6 | 0 | -0.860229 | 0.091870 | -0.000003 |
| 7 | 1 | 0 | 1.247462 | -2.582457 | 0.000007 |
| 8 | 1 | 0 | -0.180270 | 2.140615 | -0.000010 |
| 9 | 8 | 0 | 2.352253 | 1.848788 | -0.000007 |
| 10 | 1 | 0 | 3.263069 | 1.527526 | -0.000005 |
| 11 | 6 | 0 | 3.385441 | -0.884623 | 0.000005 |
| 12 | 1 | 0 | 3.895048 | -0.479990 | -0.882044 |
| 13 | 6 | 0 | -2.338388 | 0.455028 | -0.000004 |
| 14 | 1 | 0 | -2.397383 | 1.548465 | -0.000015 |
| 15 | 1 | 0 | -1.152196 | -2.049151 | 0.000002 |
| 16 | 1 | 0 | 3.895048 | -0.479977 | 0.882049 |
| 17 | 1 | 0 | 3.530828 | -1.965886 | 0.000013 |
| 18 | 6 | 0 | -3.051295 | -0.050291 | 1.266041 |
| 19 | 1 | 0 | -2.571289 | 0.331770 | 2.171582 |
| 20 | 1 | 0 | -4.095248 | 0.277940 | 1.272194 |
| 21 | 1 | 0 | -3.043914 | -1.143485 | 1.314632 |
| 22 | 6 | 0 | -3.051303 | -0.050318 | -1.266033 |
| 23 | 1 | 0 | -4.095256 | 0.277912 | -1.272186 |
| 24 | 1 | 0 | -2.571304 | 0.331724 | -2.171585 |
| 25 | 1 | 0 | -3.043922 | -1.143513 | -1.314601 |

Rotational constants (GHZ): 2.2667439 0.6387861 0.5605769

**Bromothymol**

Free Energy -3038.245355
Stoichiometry C10H13BrO
Framework group C1[X(C10H13BrO)]
Deg. of freedom 69
Full point group C1 NOp 1
Largest Abelian subgroup C1 NOp 1
Largest concise Abelian subgroup C1 NOp 1
Standard orientation:

| Center Number | Atomic Number | Atomic Type | Coordinates (Angstroms) X | Y | Z |
|---|---|---|---|---|---|
| 1 | 6 | 0 | -0.926816 | 0.040259 | -0.018848 |
| 2 | 6 | 0 | 0.267779 | -0.672063 | -0.080228 |
| 3 | 6 | 0 | 1.501466 | -0.024683 | -0.147859 |
| 4 | 6 | 0 | 1.480644 | 1.379047 | -0.150045 |
| 5 | 6 | 0 | 0.280501 | 2.083079 | -0.092832 |
| 6 | 6 | 0 | -0.958317 | 1.436176 | -0.025159 |
| 7 | 1 | 0 | 0.229983 | -1.752922 | -0.080517 |
| 8 | 1 | 0 | 0.302139 | 3.169920 | -0.101067 |
| 9 | 8 | 0 | 2.688872 | 2.028976 | -0.210775 |
| 10 | 1 | 0 | 2.543853 | 2.980269 | -0.236563 |
| 11 | 6 | 0 | 2.827558 | -0.771087 | -0.185558 |
| 12 | 1 | 0 | 3.499846 | -0.178150 | -0.813531 |
| 13 | 6 | 0 | 3.455444 | -0.833708 | 1.222148 |
| 14 | 1 | 0 | 4.438904 | -1.311957 | 1.182445 |
| 15 | 1 | 0 | 3.581870 | 0.165768 | 1.642951 |
| 16 | 1 | 0 | 2.823492 | -1.413994 | 1.901689 |
| 17 | 6 | 0 | 2.730571 | -2.172671 | -0.803009 |
| 18 | 1 | 0 | 2.268122 | -2.146860 | -1.793276 |
| 19 | 1 | 0 | 3.731037 | -2.600605 | -0.909496 |
| 20 | 1 | 0 | 2.153119 | -2.858594 | -0.175787 |
| 21 | 6 | 0 | -2.234196 | 2.233758 | 0.034376 |
| 22 | 1 | 0 | -2.022249 | 3.304180 | 0.017507 |
| 23 | 1 | 0 | -2.887293 | 1.999563 | -0.810839 |
| 24 | 1 | 0 | -2.799054 | 2.009778 | 0.943535 |

| | | | | | |
|---|---|---|---|---|---|
| 25 | 35 | 0 | -2.569216 | -0.965338 | 0.068606 |

---------------------------------------------------------------------

Rotational constants (GHZ): 1.0571415 0.4057908 0.3082648

**Thymyl acetate**

Free Energy -617.351173

Stoichiometry C12H16O2

Framework group C1[X(C12H16O2)]

Deg. of freedom 84

Full point group C1 NOp 1

Largest Abelian subgroup C1 NOp 1

Largest concise Abelian subgroup C1 NOp 1

Standard orientation:

| Center Number | Atomic Number | Atomic Type | Coordinates (Angstroms) X | Y | Z |
|---|---|---|---|---|---|
| 1 | 6 | 0 | -2.358749 | 1.167877 | 0.280467 |
| 2 | 6 | 0 | -1.078078 | 1.710855 | 0.355573 |
| 3 | 6 | 0 | 0.063162 | 0.965886 | 0.039138 |
| 4 | 6 | 0 | -0.156917 | -0.362084 | -0.349266 |
| 5 | 6 | 0 | -1.430932 | -0.910871 | -0.446104 |
| 6 | 6 | 0 | -2.559788 | -0.154201 | -0.123971 |
| 7 | 1 | 0 | -3.213856 | 1.786850 | 0.531754 |
| 8 | 1 | 0 | -0.968669 | 2.743457 | 0.664393 |
| 9 | 1 | 0 | -1.532878 | -1.933908 | -0.793160 |
| 10 | 6 | 0 | 1.467196 | 1.555784 | 0.048123 |
| 11 | 1 | 0 | 2.155657 | 0.755859 | 0.338130 |
| 12 | 6 | 0 | 1.871928 | 2.001302 | -1.372301 |
| 13 | 1 | 0 | 2.899096 | 2.377813 | -1.377035 |
| 14 | 1 | 0 | 1.811612 | 1.170606 | -2.077863 |
| 15 | 1 | 0 | 1.214847 | 2.801372 | -1.726967 |
| 16 | 6 | 0 | 1.648987 | 2.701235 | 1.054108 |
| 17 | 1 | 0 | 1.341933 | 2.409501 | 2.062225 |
| 18 | 1 | 0 | 2.701163 | 2.995084 | 1.094193 |
| 19 | 1 | 0 | 1.077595 | 3.589665 | 0.769828 |
| 20 | 6 | 0 | -3.942615 | -0.754908 | -0.196656 |
| 21 | 1 | 0 | -4.018427 | -1.485158 | -1.005805 |
| 22 | 1 | 0 | -4.197014 | -1.271291 | 0.735448 |
| 23 | 1 | 0 | -4.700429 | 0.013954 | -0.361912 |
| 24 | 8 | 0 | 0.933119 | -1.137800 | -0.757165 |
| 25 | 6 | 0 | 1.514276 | -2.081617 | 0.057640 |
| 26 | 8 | 0 | 2.406463 | -2.744132 | -0.391862 |
| 27 | 6 | 0 | 0.987115 | -2.203213 | 1.468722 |
| 28 | 1 | 0 | 0.947854 | -1.231289 | 1.964554 |
| 29 | 1 | 0 | -0.028806 | -2.604265 | 1.465278 |
| 30 | 1 | 0 | 1.640166 | -2.879059 | 2.016323 |

---------------------------------------------------------------------

Rotational constants (GHZ): 0.7442776 0.6063049 0.3903740

- **B3LYP 6-311G+(d,p) heptane**

**Thymol**

Free Energy -464.693203

Stoichiometry C10H14O

Framework group C1[X(C10H14O)]

Deg. of freedom 69

Full point group C1 NOp 1

Largest Abelian subgroup C1 NOp 1

Largest concise Abelian subgroup C1 NOp 1

Standard orientation:

| Center Number | Atomic Number | Atomic Type | Coordinates (Angstroms) X | Y | Z |
|---|---|---|---|---|---|
| 1 | 6 | 0 | -1.471101 | -1.415893 | 0.147065 |
| 2 | 6 | 0 | -0.079680 | -1.340661 | 0.057090 |
| 3 | 6 | 0 | 0.591968 | -0.130216 | -0.116975 |
| 4 | 6 | 0 | -0.206075 | 1.025650 | -0.197940 |
| 5 | 6 | 0 | -1.594994 | 0.959516 | -0.113137 |
| 6 | 6 | 0 | -2.251865 | -0.263329 | 0.061242 |
| 7 | 1 | 0 | -1.947941 | -2.380969 | 0.281515 |

| | | | | | |
|---|---|---|---|---|---|
| 8 | 1 | 0 | 0.493939 | -2.257986 | 0.121259 |
| 9 | 1 | 0 | -2.173998 | 1.877955 | -0.183747 |
| 10 | 8 | 0 | 0.438314 | 2.227067 | -0.364130 |
| 11 | 1 | 0 | -0.213120 | 2.934431 | -0.429389 |
| 12 | 6 | 0 | 2.109248 | -0.016160 | -0.185111 |
| 13 | 1 | 0 | 2.333328 | 0.806499 | -0.871130 |
| 14 | 6 | 0 | 2.690585 | 0.368929 | 1.190390 |
| 15 | 1 | 0 | 3.773307 | 0.517975 | 1.124096 |
| 16 | 1 | 0 | 2.246305 | 1.293937 | 1.564989 |
| 17 | 1 | 0 | 2.502690 | -0.419940 | 1.926155 |
| 18 | 6 | 0 | 2.799464 | -1.273225 | -0.731174 |
| 19 | 1 | 0 | 2.390425 | -1.570551 | -1.700992 |
| 20 | 1 | 0 | 3.868656 | -1.082710 | -0.863204 |
| 21 | 1 | 0 | 2.706335 | -2.123855 | -0.049012 |
| 22 | 6 | 0 | -3.757887 | -0.318105 | 0.140753 |
| 23 | 1 | 0 | -4.142049 | 0.352132 | 0.916179 |
| 24 | 1 | 0 | -4.217176 | -0.014621 | -0.806144 |
| 25 | 1 | 0 | -4.105200 | -1.327862 | 0.369241 |

---------------------------------------------------------------------

Rotational constants (GHZ): 2.0296495 0.7317527 0.5894721

**Carvacrol**

Free Energy -464.694444

Stoichiometry C10H14O

Framework group C1[X(C10H14O)]

Deg. of freedom 69

Full point group C1 NOp 1

Largest Abelian subgroup C1 NOp 1

Largest concise Abelian subgroup C1 NOp 1

Standard orientation:

---------------------------------------------------------------------

| Center Number | Atomic Number | Atomic Type | Coordinates (Angstroms) X | Y | Z |
|---|---|---|---|---|---|
| 1 | 6 | 0 | 0.425454 | -1.242954 | 0.000016 |
| 2 | 6 | 0 | -0.932467 | -1.543788 | 0.000055 |
| 3 | 6 | 0 | -1.911734 | -0.546040 | 0.000030 |
| 4 | 6 | 0 | -1.470216 | 0.784019 | -0.000010 |
| 5 | 6 | 0 | -0.110229 | 1.093651 | -0.000061 |
| 6 | 6 | 0 | 0.857192 | 0.089722 | -0.000055 |
| 7 | 1 | 0 | -1.248240 | -2.582513 | 0.000093 |
| 8 | 1 | 0 | 0.175870 | 2.139865 | -0.000083 |
| 9 | 8 | 0 | -2.342399 | 1.844521 | -0.000068 |
| 10 | 1 | 0 | -3.249677 | 1.523878 | 0.000095 |
| 11 | 6 | 0 | -3.384246 | -0.877255 | 0.000003 |
| 12 | 1 | 0 | -3.897102 | -0.481176 | 0.885443 |
| 13 | 6 | 0 | 2.336045 | 0.450595 | -0.000040 |
| 14 | 1 | 0 | 2.395715 | 1.544772 | -0.000136 |
| 15 | 1 | 0 | 1.146633 | -2.052344 | 0.000036 |
| 16 | 1 | 0 | -3.897044 | -0.481154 | -0.885446 |
| 17 | 1 | 0 | -3.536282 | -1.957589 | 0.000006 |
| 18 | 6 | 0 | 3.049239 | -0.049652 | -1.269331 |
| 19 | 1 | 0 | 2.563639 | 0.331423 | -2.171295 |
| 20 | 1 | 0 | 4.093007 | 0.278895 | -1.278899 |
| 21 | 1 | 0 | 3.043902 | -1.142340 | -1.323411 |
| 22 | 6 | 0 | 3.049090 | -0.049418 | 1.269416 |
| 23 | 1 | 0 | 4.092866 | 0.279107 | 1.279048 |
| 24 | 1 | 0 | 2.563392 | 0.331822 | 2.171260 |
| 25 | 1 | 0 | 3.043737 | -1.142100 | 1.323701 |

---------------------------------------------------------------------

Rotational constants (GHZ): 2.2686748 0.6399020 0.5619228

**Bromothymol**

Free Energy -3038.249623

Stoichiometry C10H13BrO

Framework group C1[X(C10H13BrO)]

Deg. of freedom 69

Full point group C1 NOp 1

Largest Abelian subgroup C1 NOp 1

Largest concise Abelian subgroup C1 NOp 1

Standard orientation:

---------------------------------------------------------------------

| Center Number | Atomic Number | Atomic Type | Coordinates (Angstroms) X | Y | Z |
|---|---|---|---|---|---|
| 1 | 6 | 0 | -0.926816 | 0.040259 | -0.018848 |
| 2 | 6 | 0 | 0.267779 | -0.672063 | -0.080228 |
| 3 | 6 | 0 | 1.501466 | -0.024683 | -0.147859 |
| 4 | 6 | 0 | 1.480644 | 1.379047 | -0.150045 |
| 5 | 6 | 0 | 0.280501 | 2.083079 | -0.092832 |
| 6 | 6 | 0 | -0.958317 | 1.436176 | -0.025159 |
| 7 | 1 | 0 | 0.229983 | -1.752922 | -0.080517 |
| 8 | 1 | 0 | 0.302139 | 3.169920 | -0.101067 |
| 9 | 8 | 0 | 2.688872 | 2.028976 | -0.210775 |
| 10 | 1 | 0 | 2.543853 | 2.980269 | -0.236563 |
| 11 | 6 | 0 | 2.827558 | -0.771087 | -0.185558 |
| 12 | 1 | 0 | 3.499846 | -0.178150 | -0.813531 |
| 13 | 6 | 0 | 3.455444 | -0.833708 | 1.222148 |
| 14 | 1 | 0 | 4.438904 | -1.311957 | 1.182445 |
| 15 | 1 | 0 | 3.581870 | 0.165768 | 1.642951 |
| 16 | 1 | 0 | 2.823492 | -1.413994 | 1.901689 |
| 17 | 6 | 0 | 2.730571 | -2.172671 | -0.803009 |
| 18 | 1 | 0 | 2.268122 | -2.146860 | -1.793276 |
| 19 | 1 | 0 | 3.731037 | -2.600605 | -0.909496 |
| 20 | 1 | 0 | 2.153119 | -2.858594 | -0.175787 |
| 21 | 6 | 0 | -2.234196 | 2.233758 | 0.034376 |
| 22 | 1 | 0 | -2.022249 | 3.304180 | 0.017507 |
| 23 | 1 | 0 | -2.887293 | 1.999563 | -0.810839 |
| 24 | 1 | 0 | -2.799054 | 2.009778 | 0.943535 |
| 25 | 35 | 0 | -2.569216 | -0.965338 | 0.068606 |

Rotational constants (GHZ): 1.0571415 0.4057908 0.3082648

**Thymyl acetate**

Free Energy -617.354482

Stoichiometry C12H16O2

Framework group C1[X(C12H16O2)]

Deg. of freedom 84

Full point group C1 NOp 1

Largest Abelian subgroup C1 NOp 1

Largest concise Abelian subgroup C1 NOp 1

Standard orientation:

| Center Number | Atomic Number | Atomic Type | Coordinates (Angstroms) X | Y | Z |
|---|---|---|---|---|---|
| 1 | 6 | 0 | -2.358749 | 1.167877 | 0.280467 |
| 2 | 6 | 0 | -1.078078 | 1.710855 | 0.355573 |
| 3 | 6 | 0 | 0.063162 | 0.965886 | 0.039138 |
| 4 | 6 | 0 | -0.156917 | -0.362084 | -0.349266 |
| 5 | 6 | 0 | -1.430932 | -0.910871 | -0.446104 |
| 6 | 6 | 0 | -2.559788 | -0.154201 | -0.123971 |
| 7 | 1 | 0 | -3.213856 | 1.786850 | 0.531754 |
| 8 | 1 | 0 | -0.968669 | 2.743457 | 0.664393 |
| 9 | 1 | 0 | -1.532878 | -1.933908 | -0.793160 |
| 10 | 6 | 0 | 1.467196 | 1.555784 | 0.048123 |
| 11 | 1 | 0 | 2.155657 | 0.755859 | 0.338130 |
| 12 | 6 | 0 | 1.871928 | 2.001302 | -1.372301 |
| 13 | 1 | 0 | 2.899096 | 2.377813 | -1.377035 |
| 14 | 1 | 0 | 1.811612 | 1.170606 | -2.077863 |
| 15 | 1 | 0 | 1.214847 | 2.801372 | -1.726967 |
| 16 | 6 | 0 | 1.648987 | 2.701235 | 1.054108 |
| 17 | 1 | 0 | 1.341933 | 2.409501 | 2.062225 |
| 18 | 1 | 0 | 2.701163 | 2.995084 | 1.094193 |
| 19 | 1 | 0 | 1.077595 | 3.589665 | 0.769828 |
| 20 | 6 | 0 | -3.942615 | -0.754908 | -0.196656 |
| 21 | 1 | 0 | -4.018427 | -1.485158 | -1.005805 |
| 22 | 1 | 0 | -4.197014 | -1.271291 | 0.735448 |
| 23 | 1 | 0 | -4.700429 | 0.013954 | -0.361912 |
| 24 | 8 | 0 | 0.933119 | -1.137800 | -0.757165 |
| 25 | 6 | 0 | 1.514276 | -2.081617 | 0.057640 |
| 26 | 8 | 0 | 2.406463 | -2.744132 | -0.391862 |
| 27 | 6 | 0 | 0.987115 | -2.203213 | 1.468722 |
| 28 | 1 | 0 | 0.947854 | -1.231289 | 1.964554 |

| | | | | | |
|---|---|---|---|---|---|
| 29 | 1 | 0 | -0.028806 | -2.604265 | 1.465278 |
| 30 | 1 | 0 | 1.640166 | -2.879059 | 2.016323 |

---------------------------------------------------------------------

Rotational constants (GHZ): 0.7442776 0.6063049 0.3903740
